# MAR1 links membrane adhesion to membrane merger during cell-cell fusion in *Chlamydomonas*

**DOI:** 10.1101/2021.09.03.458930

**Authors:** Jennifer F. Pinello, Yanjie Liu, William J. Snell

## Abstract

Union of two gametes to form a zygote is a defining event in the life of sexual eukaryotes, yet the mechanisms that underlie cell-cell fusion during fertilization remain poorly characterized. Here, in studies of fertilization in the green alga, *Chlamydomonas*, we report identification of a membrane protein on *minus* gametes, <u>M</u>inus <u>A</u>dhesion <u>R</u>eceptor <u>1</u> (MAR1), that is essential for the membrane attachment with *plus* gametes that immediately precedes lipid bilayer merger. We show that MAR1 forms a receptor pair with previously identified receptor FUS1 on *plus* gametes, whose ectodomain architecture we find is identical to a sperm adhesion protein conserved throughout plant lineages. Strikingly, before fusion, MAR1 is biochemically and functionally associated with the ancient, evolutionarily conserved eukaryotic class II fusion protein HAP2 on *minus* gametes. Thus, the integral membrane protein MAR1 provides a molecular link between membrane adhesion and bilayer merger during fertilization in *Chlamydomonas*.

## Introduction

Fusion of gametes to form a zygote during fertilization is a complex cell-cell interaction whose core cellular features have been conserved since the origin of eukaryotes. Initial cell-cell recognition, typically at cell surface locations entirely separate from the site of fusion, bring the two gametes together and induce signals within them to prepare for fusion (Dean, 2007; Dresselhaus et al., 2016; Ikawa et al., 2010; Snell and Goodenough, 2009). Upon the consequent gamete activation, the gametes undergo a second interaction by gamete-specific adhesion proteins at newly available sites on their plasma membranes specialized for fusion. Attachment of the plasma membranes is rapidly followed by merger of the lipid bilayers to complete the fusion reaction. The conservation of the cellular steps of fertilization suggests that at least some of the molecular underpinnings of the gamete membrane fusion reaction might also be conserved (Sankaranarayanan and Higashiyama, 2018; Tajima and Nishimura, 2018). Surprisingly, in spite of over a century of study of fertilization (Bianchi and Wright, 2020; Lillie, 1914), we still lack a comprehensive understanding in any single organism of the molecules or mechanisms that compose and regulate the gamete fusion reaction.

Studies in mouse have shown that the proteins IZUMO1 on sperm and JUNO on the egg form a complex that is essential for the sperm-egg membrane binding preceding fusion. The two proteins, which form the only receptor pair known to be essential for gamete membrane adhesion in any organism, are insufficient for fusion and a chordate gamete fusogen remains unidentified (Aydin et al., 2016; Bianchi et al., 2014; Inoue et al., 2005; Ohto et al., 2016). Several other mouse gamete proteins with mammalian orthologs have been reported as essential for fertilization, including CD9 on eggs (Le Naour et al., 2000; Miyado et al., 2000), and FIMP (Fujihara et al., 2020), SPACA6 (Barbaux et al., 2020; Lorenzetti et al., 2014), SOF1 (Noda et al., 2020), and TMEM95 (Lamas-Toranzo et al., 2020; Noda *et al.*, 2020) on sperm, but they neither adhere nor fuse cells when heterologously expressed, and their molecular functions remain unclear.

Proteins with roles in gamete membrane interactions during the fusion reaction have been described in several other model organisms, including Prm1p in *S. cerevisiae* (Heiman and Walter, 2000); Spe-9, Spe-45, and EGG-1 & 2 in *C. elegans* (Kadandale et al., 2005; Nishimura et al., 2015; Singaravelu et al., 2015; Singson et al., 1998); P48/45, p47 and p230 in the malaria pathogens *Plasmodium* (van Dijk et al., 2010); the mating type proteins in ciliates (Cervantes et al., 2013); Bouncer in *D. rerio* (Herberg et al., 2018); GEX2 and DMP8 & 9 in the plant *Arabidopsis thaliana* (Cyprys et al., 2019; Engel et al., 2005; Mori et al., 2014; Takahashi et al., 2018); and FUS1 in the green alga *Chlamydomonas reinhardtii* (hereafter, *Chlamydomonas*) (Ferris et al., 1996; Misamore et al., 2003). The presence of immunoglobulin-like (Ig-like) domains in Spe-45, IZUMO1, GEX2, and FUS1 is consistent with the known roles of the domain in protein-protein interactions. This collection of adhesion proteins, however, lacks a common overall domain architecture, and thus, a gamete adhesion protein family that spans unicellular and multicellular plants or animals has not been reported.

In contrast, a single protein of conserved structure, HAP2, is essential for fertilization in unicellular and multicellular organisms across kingdoms, and was likely the primordial sexual fusogen (Camacho-Nuez et al., 2017; Cole et al., 2014; Ebchuqin et al., 2014; Hirai et al., 2008; Johnson et al., 2004; Liu et al., 2008; Mori et al., 2006; Okamoto et al., 2016; Ramakrishnan et al., 2019; Steele and Dana, 2009). Studies in *Chlamydomonas, Plasmodium,* and *Tetrahymena* showed that HAP2 was dispensable for gamete membrane adhesion, but was required for bilayer merger (Cole *et al.*, 2014; Liu *et al.*, 2008). Recent studies uncovered an unambiguous structural homology between HAP2 and viral and developmental class II fusion proteins, including the envelope proteins from the *Bunyavirales*, *Flaviviruses,* and *Alphaviruses* and EFF-1 from *C. elegans* (Fedry et al., 2017; Perez-Vargas et al., 2014; Pinello et al., 2017; Valansi et al., 2017). Notably, HAP2 (also called GCS1) has not yet been detected in fungi or chordates, either because it was lost or because it evolved to become unrecognizable.

In *Chlamydomonas*, when *plus* and *minus* gametes are mixed together, they initially recognize and adhere to each other by their cilia through mating type-specific adhesion proteins that function only on the cilia, SAG1 of *plus* gametes and SAD1 of *minus* gametes (Snell and Goodenough, 2009) (Fig. 1A). Ciliary adhesion activates a protein kinase-dependent signaling pathway that leads to increases in intracellular cyclic AMP, thereby triggering gamete activation. During activation, gametes release their extracellular matrices (cell walls), recruit additional adhesion proteins to their ciliary membranes (Belzile et al., 2013; Ranjan et al., 2019), and assemble a membrane protuberance called a mating structure between their two cilia (Friedmann et al., 1968; Weiss et al., 1977). Ciliary adhesion and the accompanying vigorous motility of the cells bring the apical ends of a pair of gametes into alignment, leading to collisions between the two mating structures and adhesion of their tips. FUS1 is the membrane adhesion protein on *plus* gamete mating structures, and *plus* gametes lacking FUS1 fail to adhere and fail to fuse (Ferris *et al.*, 1996; Misamore *et al.*, 2003). Within seconds of membrane adhesion, HAP2 on the *minus* mating structure engages with the membrane of the *plus* mating structure to fuse the two membranes, followed by rapid coalescence of the two gametes into a quadri-ciliated zygote (Fig. 1A). Nuclear fusion by the conserved KAR5 family member, GEX1, follows soon thereafter (Beh et al., 1997; Engel *et al.*, 2005; Ning et al., 2013).

**Figure 1.**
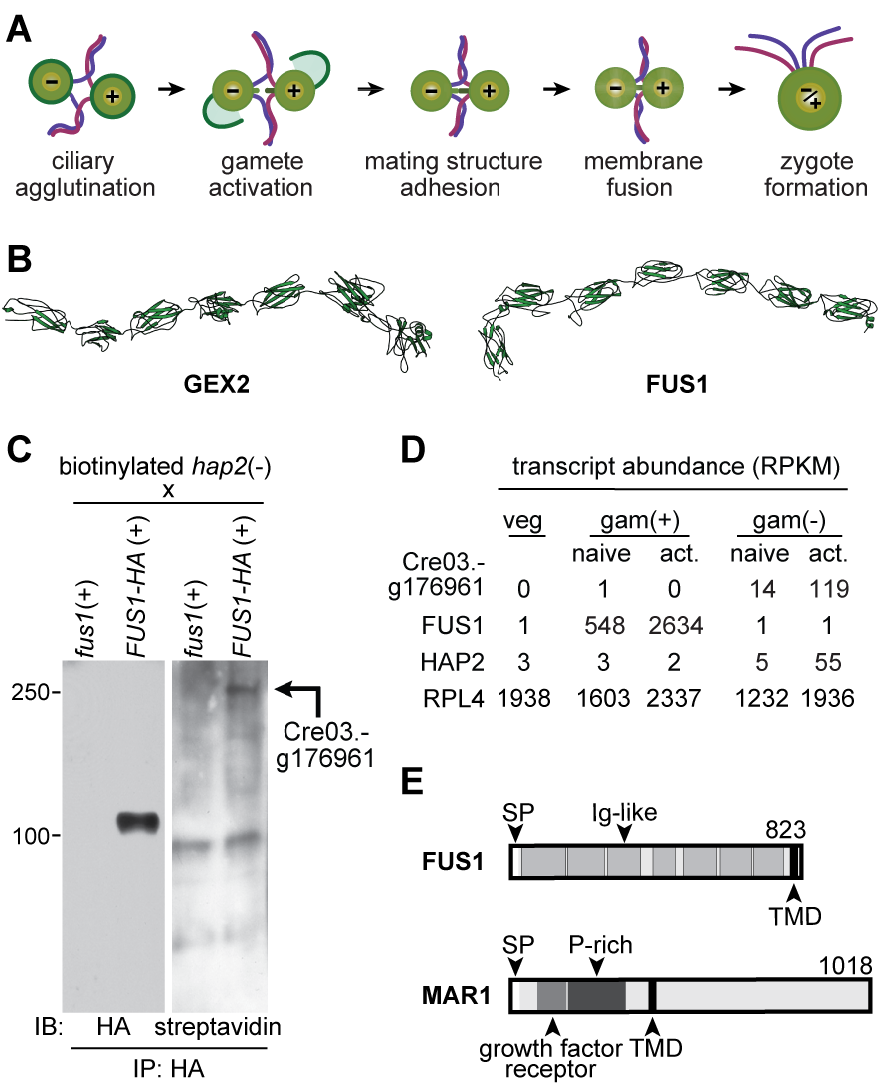
Identification of a FUS1 binding protein in *minus* gametes. (A) Illustration depicting steps (left to right) in *Chlamydomonas* fertilization. **(B)** RaptorX structural models of *A. thaliana* GEX2 and *C. reinhardtii* FUS1 ectodomains. Each model is orientated N’ terminus (left) to C’ terminus (right). **(C)** The Cre03.g176961 gene product is a FUS1-HA binding protein on *minus* gametes. Immunoblots (IB) probed with anti-HA antibodies (left) or streptavidin-HRP (right) showing anti-HA immunoprecipitates (IP) of lysate supernatants from live, biotinylated *hap2(-)* gametes that had been mixed with *fus1(+)* or *FUS1-HA(+)* gametes. The identity of the biotinylated protein detected in the FUS1-HA(+) immunoprecipitates, Cre03.g176961, was found by mass spectrometry. **(D)** Cre03.g176961 transcripts are specifically expressed in *minus* gametes. RPKM (Reads Per Kilobase per Million mapped reads) data from Ning et al. (2013) of the indicated transcripts (left) in vegetative cells (veg) and in naive and activated *plus* and *minus* gametes; gam(+) and gam(-). A constitutively expressed transcript encoding ribosomal protein, RPL4 (Cre09.g397697) is also shown. **(E)** Domain architecture illustrations of FUS1 (top) and MAR1 (bottom) proteins, showing the signal peptide (SP) transmembrane domain (TMD) in each, the Ig-like folds (Ig-like) in FUS1, and the growth factor receptor cysteine-rich domain and proline-rich region (P-rich) in MAR1.

Here, we report that *Chlamydomonas* FUS1 and previously described plant sperm adhesion protein, GEX2, are members of a broadly conserved protein family characterized by extracellular domains composed entirely of Ig-like domains. We identify a new, lineage-restricted protein on *minus* gametes, MAR1, that is a receptor for FUS1 on *plus* gametes. Formation of the FUS1-MAR1 pair is essential for the gamete membrane attachment that immediately precedes bilayer merger and is necessary for gamete fusion. Moreover, we find that MAR1 is biochemically associated with HAP2 and is required for proper HAP2 expression and localization at the *minus* mating structure. Thus, during the gamete membrane fusion reaction in *Chlamydomonas*, a lineage-specific protein, MAR1, functions at the nexus of conserved membrane adhesion protein, FUS1, and ancient membrane fusogen, HAP2.

## Results

### *Chlamydomonas* FUS1 and plant gamete adhesion protein, GEX2, share a common ectodomain architecture

We used the more powerful protein analysis methods available now to update the earlier reports that the FUS1 ectodomain contained 5 Ig-like domains (Misamore *et al.*, 2003). Our new analysis uncovered FUS1 orthologs in other algae species (Supplemental Fig. S3) and identified two more Ig-like domains in FUS1 (for a total of 7). To our surprise, DELTA-BLAST and PSI-BLAST searches identified sequence similarity between the ectodomains of *Chlamydomonas* FUS1 and several plant GEX2 sequences (including those of *Gossypium* sp., *Durio zibethinus*, *Theobromoa cacao*, *Hibiscus syriacus*, *Vigna angularis*). A PROMALS3D (Pei and Grishin, 2007) pairwise alignment confirmed and extended the BLAST analyses, showing similarities in sequence and secondary structure between the entire ectodomains of *Chlamydomonas* FUS1 and *Arabidopsis* GEX2 (∼16% sequence identity, Supplemental Fig. S4). Consistent with these results, RaptorX structural models (Xu, 2019) showed similar tertiary structures for *Chlamydomonas* FUS1 and *Arabidopsis* GEX2 ectodomains, with both predicted to be composed entirely of 7 linearly-arranged Ig-like domains (Fig. 1B, E). Furthermore, additional structural homology modeling on all platforms tested (PHYRE2, SWISS-MODEL and HHPRED) found that *Chlamydomonas* FUS1 and *Arabidopsis* GEX2 have strong homologies to the same template structures in the protein data bank (Supplemental Fig. S3), namely the bacterial invasin-, intimin-like proteins (e.g. PDB ID: 4E9L) and the *Dictyostelium* gelation factor rod domain (e.g. PDB ID: 1WLH), demonstrating a structural uniformity among their Ig-like domains. Taken together, our results indicate that FUS1 and GEX2 belong to a family of gamete cell surface adhesion proteins whose ectodomains are constituted by bacterial-type Ig-like domains.

### A *minus* gamete protein that binds to FUS1

We took advantage of the ease of preparing biochemical quantities of pure gametes in *Chlamydomonas* to identify proteins on *minus* [*(-)*] gametes that bound to FUS1 on *plus* [*(+)*] gametes. *hap2(-)* gametes were surface labeled with biotin, mixed with *fus1*::*FUS1-HA(+)* gametes, and immunoprecipitation, immunoblotting, and mass spectrometry were used to identify biotinylated *minus* gamete proteins that associated with FUS1-HA. Use of the fusion-defective *hap2(-)* gametes to block fertilization at the stage of mating structure adhesion maximized opportunities for adhesion protein interactions and prevented the degradation of FUS1 that is coincident with gamete fusion (Liu et al., 2010). As a control, we also mixed biotinylated *hap2(-)* gametes with *fus1(+)* gametes, which lack FUS1. FUS1-HA was pulled down efficiently by anti-HA antibody, and as expected, staining was absent in the *fus1* control samples (Fig. 1C, left panel). Importantly, streptavidin blotting showed a biotinylated protein migrating with an apparent molecular mass of ∼250 kDa that was co-immunoprecipitated only in the FUS1-HA-containing sample (Fig. 1C, right panel).

### Specific expression of Cre03.g176961 in *minus* gametes

The protein encoded by gene Cre03.g176961 exhibited high coverage in mass spectrometry analysis of the 250 kDa region of samples from FUS1 immunoprecipitates and was absent in *fus1* samples (Supplemental Fig. S1). Importantly, previous gene expression studies in *Chlamydomonas* vegetative cells and naive and activated gametes (Ning *et al.*, 2013) showed that Cre03.g176961 transcripts were uniquely present in *minus* gametes and upregulated in activated *minus* gametes (Fig. 1D). This expression signature is similar to that of other fertilization-essential transcripts in *minus* gametes (Ning *et al.*, 2013), and distinct from those of *plus*-specific *FUS1* and constitutively expressed ribosomal transcript *RPL4* (Fig.1D), making the Cre03.g176961 protein, hereafter called *<u>M</u>inus* <u>A</u>dhesion <u>R</u>eceptor <u>1</u> (MAR1), a strong candidate for a FUS1 binding partner.

Based on the annotation in Phytozome (Goodstein et al., 2012), and confirmatory RT-PCR characterization of the transcript, *MAR1* (GenBank: KT288268) encodes a 1018-residue, single-pass transmembrane protein with a predicted molecular mass of 99 kDa. Analysis of the amino acid sequence predicted a signal peptide, a 365-residue ectodomain, a single transmembrane domain, and a 608-residue cytoplasmic domain (Fig. 1E). Two notable features of the ectodomain are a growth factor receptor cysteine-rich domain near the N-terminus (residues 78-159) and a proline-rich segment spanning residues 142 to 365 that includes 5 repeats of a “PPSPX” motif. The latter motif was previously identified in other *Chlamydomonas* proteins, including a cell wall protein, GP1 (Ferris et al., 2001); and the ciliary adhesion proteins, SAG1 and SAD1, which mediate initial gamete recognition (Ferris et al., 2005). In the MAR1 cytoplasmic domain, structural homology searches found sequential segments with likeness to the ABA receptor kinase from *Arabidopsis* (PDB: 5XD6, residues 512-620), and collagen (PDB: 1YGV, residues 619-1018). Detectable MAR1 homologies at the amino acid sequence level in other organisms, however, were scant, with only a small number of close algal relatives containing putative MAR1 orthologs (Supplemental Fig. S5).

We introduced a FLAG-tagged MAR1 transgene driven by its endogenous promoter into *hap2(-)* cells, and *hap2*::*HAP2-HA(-)* cells. The MAR1-FLAG protein was expressed in *minus* gametes and absent in *minus* vegetative cells (Fig. 2A). Consistent with the biotinylation studies, and in spite of a predicted molecular mass of 107 kDa, MAR1-FLAG appeared as a ∼250 kDa protein in SDS-PAGE. We are uncertain of the basis for this anomalous behavior, but the several proline-rich regions might contribute to its altered migration pattern (Ferris *et al.*, 2001). Immunoblot analyses of lysates from equal numbers of transgene-expressing gametes indicated that the three gamete-specific proteins, MAR1, HAP2 and FUS1, were expressed at similar levels (Supplemental Fig. S2).

**Figure 2.**
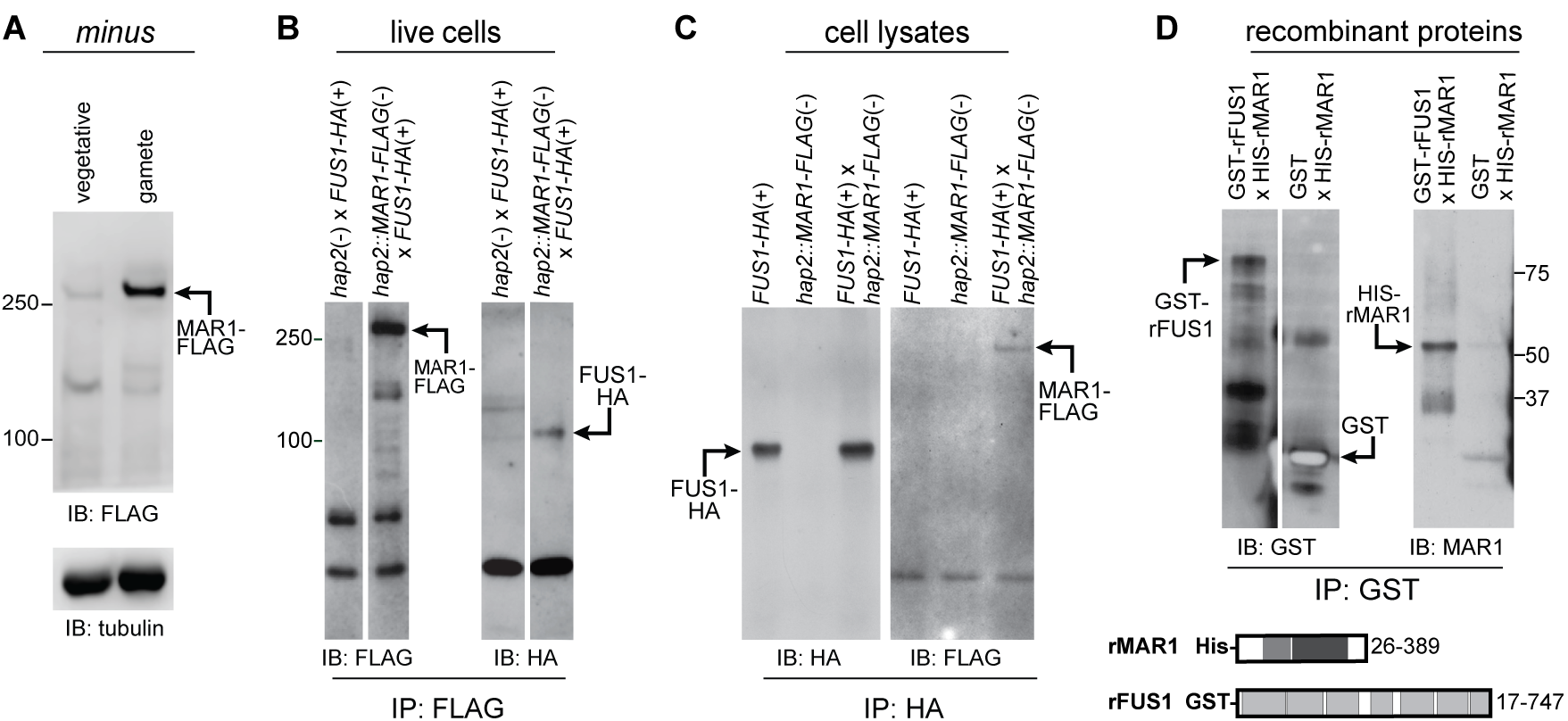
MAR1 on *minus* gametes directly interacts with FUS1 on *plus* gametes. (A) MAR1-FLAG is enriched in *minus* gametes compared to *minus* vegetative cells. Immunoblot (IB) with anti-FLAG antibodies showing equal cell equivalents of *hap2::MAR1-FLAG; HAP2-HA(-)* vegetative cells and gametes (upper panel). Lower panel shows a tubulin loading control. **(B)** MAR1-FLAG interacts with FUS1-HA during mating structure adhesion. *FUS1-HA(+)* gametes were mixed for 30 min with *hap2*::*MAR1-FLAG(-)* or *hap2(-)* gametes, and lysate supernatants from the mixed gametes were subjected to immunoprecipitation (IP) with anti-FLAG antibodies followed by immunoblotting with anti-FLAG (left) or anti-HA antibodies (right). **(C)** FUS1 and MAR1 interact with each other when separately prepared cell lysates are mixed. Separately lysed mixtures of *FUS1-HA(+)* with *hap2(-)* gametes and *hap2::MAR1-FLAG(-)* with *fus1(+)* gametes were either kept separate (controls; left and middle lanes) or mixed together (right lane) and subjected to immunoprecipitation with anti-HA antibodies followed by immunoblotting with anti-HA (left) or anti-FLAG (right) antibodies. **(D)** Recombinant GST-tagged FUS1 protein (GST-rFUS1) interacts with recombinant His-tagged MAR1 protein (His-rMAR1). GST-rFUS1 and GST protein in bacterial lysates were bound to glutathione beads followed by incubation of those beads with lysates from bacteria expressing His-rMAR1. Bound proteins eluted with reduced glutathione were immunoblotted with anti-GST (upper left) or anti-MAR1 antibodies (upper right). Illustrations depicting the ectodomain residues present in the His-rMAR1 and GST-rFUS1 recombinant proteins (lower panel).

### MAR1 is a receptor for FUS1

In vivo and in vitro protein interaction assays were consistent with the biotinylation results and showed that MAR1 and FUS1 bound to each other. In in vivo assays with live cell mixtures, *fus1::FUS1-HA(*+) gametes were mixed with *hap2::MAR1-FLAG(-)* gametes for 30 minutes to allow gamete activation and mating structure adhesion without membrane fusion, followed by disruption of the samples in lysis buffer, immunoprecipitation with anti-FLAG antibody, and immunoblotting with anti-HA antibody. *hap2(-)* gametes (i. e., expressing only un-tagged endogenous MAR1) mixed with *fus1::FUS1-HA(+)* gametes were used as a control. As shown in Fig. 2B, FUS1-HA was present in the FLAG immunoprecipitates of the *hap2::MAR1-FLAG(-)* gametes that had been mixed with *fus1::FUS1-HA(+)* gametes, but not in the control immunoprecipitates. In related experiments, separately prepared detergent lysates of activated *hap2::MAR1-FLAG(-)* gametes and *fus1::FUS1-HA(+)* gametes were subsequently mixed together followed by immunoprecipitation and immunoblotting as above, and also showed that the two proteins were associated with each other (Fig. 2C). The lysates of activated *FUS1-HA(+)* gametes and *hap2::MAR1-FLAG(-)* gametes were controls. Finally, we found that bacterially-expressed, His-tagged MAR1 ectodomain (His-MAR1) was precipitated by the GST-tagged FUS1 ectodomain (rFUS1) but not by GST alone (Fig. 2D). Together, these results demonstrated that the ectodomains of MAR1 and FUS1 directly interacted with each other.

### Mating structure adhesion is severely impaired in *mar1 minus* gametes and fusion is blocked

To test whether the MAR1 and FUS1 interaction detected biochemically was important functionally, we examined the fertilization phenotypes of a *Chlamydomonas* CLiP library mutant *minus* strain, *LMJ.RY0402.185187* (hereafter, called *mar1*), which is annotated to have an insertion in the *MAR1* gene (Li et al., 2019). PCR across the predicted insertion site showed that the *CIB1* plasmid had indeed inserted into the 3^rd^ exon of *MAR1*, thereby disrupting its coding sequence (Fig. 3A). The *mar1(-)* mutant cells showed no detectable morphological or growth phenotype when cultured as vegetative cells. Furthermore, when placed into N-free medium overnight, the *mar1(-)* cells differentiated into *minus* gametes that were fully capable of ciliary adhesion with wild type (WT) *plus* gametes upon mixing, forming characteristic, large clusters of *plus* and *minus* gametes adhering to each other by their cilia (Fig. 3B low magnification DIC images; Fig. 3C SEM image). Gamete activation, as quantitatively assessed by cell wall loss, was also indistinguishable in mixtures of *mar1(-)* gametes or WT*(-)* control gametes with WT*(+)* gametes (Fig. 3B, bottom).

**Figure 3.**
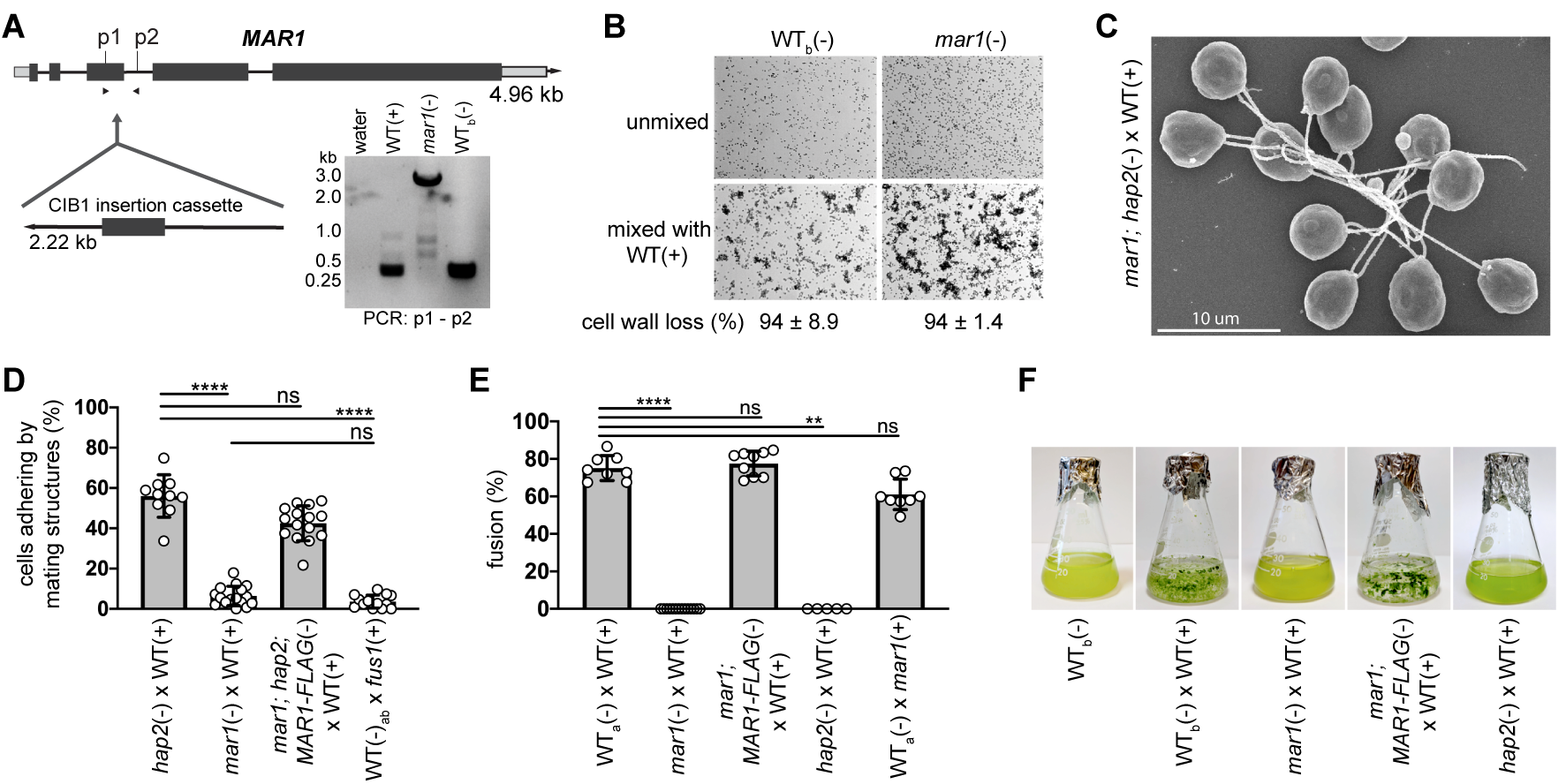
MAR1 is essential for mating structure adhesion and gamete fusion. (A) Illustration of the *MAR1* gene showing the location of the disrupting CIB1 insertion cassette in the *mar1(-)* mutant (CLiP: *LMJ.RY0402.185187*). Dark grey rectangles and lines represent exons and introns, respectively. Light grey rectangles represent 5’ and 3’ UTRs. DNA gel electrophoresis image (lower right) shows results of genotyping PCRs across the *mar1* insertion site using primers p1 and p2 and DNA template from WT*(+)*, *mar1(-)*, and WT*(-)* cells, and a water template control. The 2.6 kB amplicons from *mar1*(-) cells were sequenced to confirm the CIB1 insertion location in the 3^rd^ exon of the *MAR1* coding sequence (Supplemental Fig. S7). **(B)** Ciliary agglutination and gamete activation are unperturbed in the *mar1(-)* gametes. Low magnification differential interference contrast (DIC) microscopy images of live, WT_b_*(-)* and *mar1(-)* gametes before (top) and 10 min after (bottom) mixing with WT*(+)* gametes. The large clusters of cells visible in the bottom images form when multiple cells interact with each other by their cilia. The percent of cells that lost their cell walls upon mixing is indicated below the images (± SD). **(C)** Scanning electron micrograph of *mar1; hap2(-)* gametes experiencing ciliary adhesion with WT*(+)* gametes. **(D)** *mar1(-)* gametes fail to undergo mating structure adhesion with WT*(+)* gametes. Results are from 10 to 16 biological replicates per group analyzed using one-way non-parametric ANOVA with Dunn’s post-test; \*\*\*\**P*<0.0001; ns, not significant; bars are means; error bars, SD. **(E-F)** MAR1 is required for gamete fusion. Gamete fusion and zygote formation were assessed microscopically at 10 min **(E)** and macroscopically at 1-3 days **(F)** after mixing of the indicated gametes. For **(E),** results from 5 to 14 biological replicates per group were analyzed using one-way non-parametric ANOVA with Dunn’s post-test; \*\*\*\**P*<0.0001 and \*\**P*=0.0027; bars are means; error bars, SD. Images in **(F)** are representative of one to three biological replicates. *21gr* was the WT*(+)* gamete strain used as controls in these experiments. WT*(-)* gametes used as controls were *hap2::HAP2-HA(-)* (WT_a_) and parental CLiP mutant strain, *CC-5325(-)* (WT_b_).

*mar1(-)* gametes, however, were severely impaired in their ability to undergo mating structure adhesion with WT*(+)* gametes. Whereas a mean of 56% of cells adhered by their mating structures in the fusion-blocked mixtures of *hap2(-)* gametes with WT*(+)* gametes, only 6% adhered in the mixtures of *mar1(-)* gametes with WT*(+)* gametes. This adhesion-defective phenotype was indistinguishable from that observed in mixtures of WT*(-)* gametes mixed with *fus1(+)* gametes (Fig. 3D and Misamore *et al.*, 2003). Introduction of the *MAR1-FLAG* transgene into *mar1(-)* gametes also harboring the *hap2* gene-disruption, rescued mating structure adhesion with WT*(+)* gametes to levels analogous to that of mixtures of *hap2(-)* and WT*(+)* gametes. These results demonstrated a central role for MAR1 in mating structure adhesion.

Notably, and mirroring the phenotypes observed in both the adhesion-defective *fus1(+)* gametes (Ferris *et al.*, 1996; Misamore *et al.*, 2003) and fusion-defective *hap2(-)* gametes, loss of MAR1 completely abrogated gamete fusion as quantified by counts of quadri-ciliated zygotes at 10-30 min after mixing (Fig. 3E, 10 min results) and by visual examination after overnight incubation to detect appearance of immotile zygotes, which appear as heterogeneous, green flocculant material in liquid cultures (Fig. 3F). The fusion defect of *mar1(-)* gametes was rescued with the *MAR1-FLAG* transgene (Fig. 3E, F), and the extent of fusion 10 minutes after mixing *mar1;MAR1-FLAG*(-) gametes with WT*(+)* gametes (77% fusion) was indistinguishable from that in mixtures of WT gametes (75% fusion), consistent with the results of the macroscopic assay (Fig. 3F). As expected, and demonstrating that MAR1 functions only in *minus* gametes during fertilization, *plus* gametes bearing the *mar1* mutation were fully competent to fuse with WT*(-)* gametes (61%). These results showed that MAR1 was essential in *minus* gametes for zygote formation.

Furthermore, analysis of F_1_ and F_2_ progeny from crosses of *MAR1-FLAG*-rescued *mar1(-)* gametes mixed with *plus* gametes (Supplemental Fig. S7) confirmed that the adhesion- and fusion-defective phenotypes of *mar1(-)* gametes co-segregated with the *mar1* mutant genotype. As expected, crosses of *mar1(-)* gametes with WT*(+)* gametes failed to produce any progeny. Taken together, our biochemical and genetic results demonstrated that the direct interaction of mating structure membrane proteins MAR1 on *minus* gametes and FUS1 on *plus* gametes mediated the membrane attachment between the two gametes that occurs after their mutual recognition and activation, and that formation of this receptor pair was essential for the gamete membrane fusion reaction.

### MAR1-FLAG is localized at the *minus* mating structure and is lost rapidly after gamete fusion

Confirming that MAR1 functions at the cell surface, brief incubation of live *hap2::MAR1-FLAG; HAP2-HA(-)* gametes with pronase resulted in loss of nearly all of MAR1-FLAG (Fig. 4A, top), and, as expected, only the surface-expressed upper isoform of HAP2 (Fig. 4A, middle, Liu *et al.*, 2008). Immunostaining of *hap2::MAR1-FLAG; HAP2-HA(-)* gametes with anti-FLAG antibodies further showed that MAR1-FLAG was present as a punctum between the two cilia at the apical ends of the *minus* gametes (Fig. 4B), the location of the *minus* mating structure. Double immunostaining of *hap2::MAR1-FLAG; HAP2-HA(-)* gametes with anti-HA and anti-FLAG antibodies further confirmed their co-localization at the *minus* mating structure (Fig. 4C). Finally, similar to loss of HAP2 and FUS1 after fusion, and as part of a block to polygamy (Liu *et al.*, 2010), 30 minutes after ∼70% of *mar1::MAR1-FLAG(-)* gametes had fused with WT*(+)* gametes, MAR1-FLAG had become nearly undetectable (Fig. 4D).

**Figure 4.**
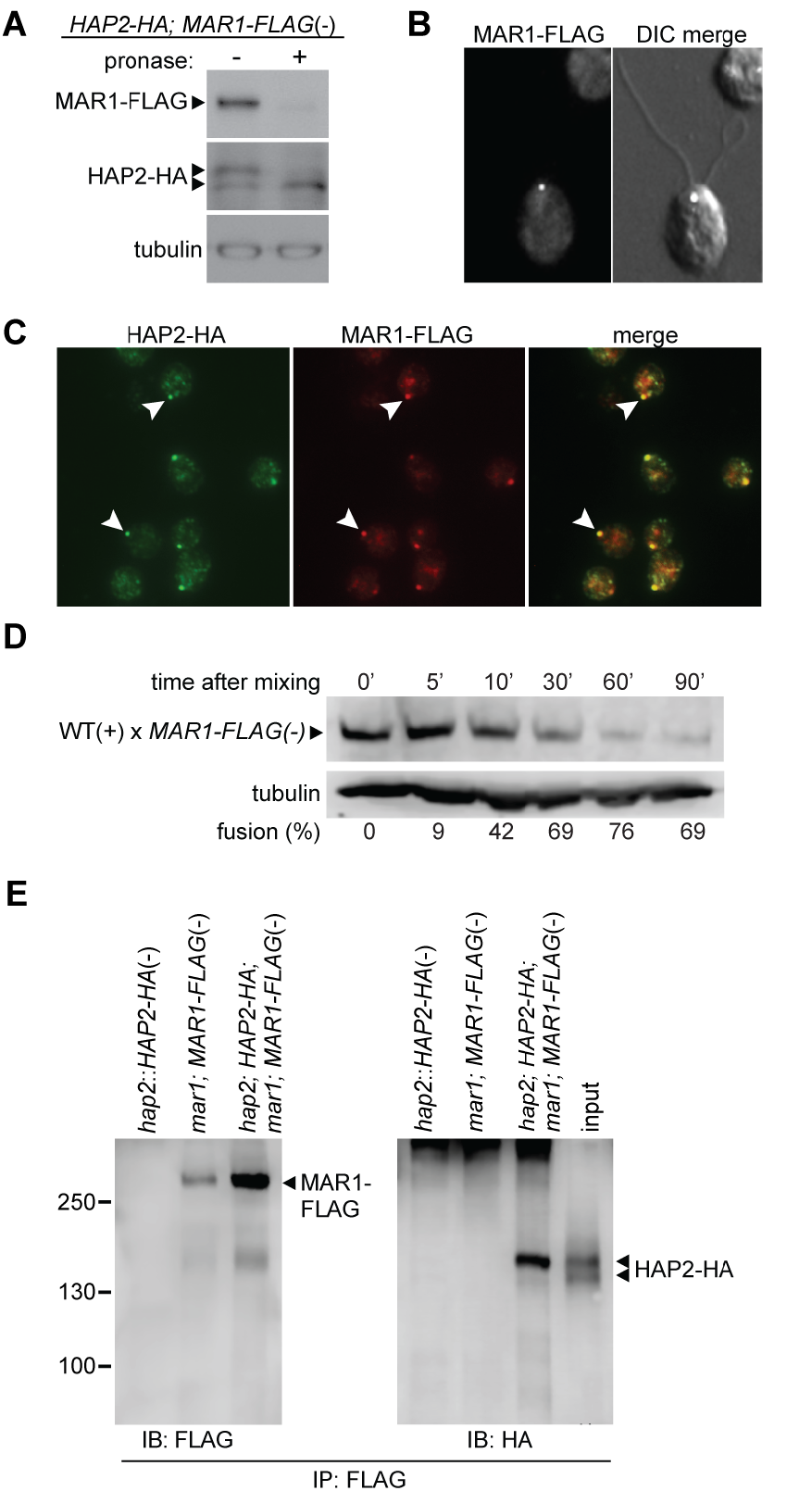
MAR1 co-localizes with HAP2 on the surface of the *minus* mating structure and is biochemically associated with the fusogen. (A) MAR1-FLAG is expressed at the cell surface. Live *hap2::HAP2-HA; MAR1-FLAG(-)* gametes were incubated with or without pronase (0.05%) for 20 min followed by immunoblotting with anti-FLAG and anti-HA antibodies. **(B)** MAR1-FLAG is localized at the *minus* mating structure at the apical end of *minus* gametes between the two cilia. Merged DIC and fluorescence images of a naive *hap2::HAP2-HA; MAR1-FLAG(-)* gamete immunostained with anti-FLAG antibodies. **(C)** MAR1-FLAG staining is coincident with HAP2-HA at the *minus* gamete mating structure (arrowheads). *hap2::MAR1-FLAG; HAP2-HA(-)* gametes were immunostained with anti-FLAG and anti-HA antibodies **(D)** MAR1-FLAG is rapidly lost after gamete fusion. *mar1::MAR1-FLAG(-)* gametes were mixed with WT*(+)* gametes and assayed for fusion and for the presence of MAR1-FLAG at the indicated times after mixing. The lower panel shows tubulin loading controls. Percent fusion is shown below the blots. **(E)** MAR1 is biochemically associated with HAP2. *Minus* gametes expressing both MAR1-FLAG and HAP2-HA were activated by mixing with adhesion-defective *fus1(+)* gametes for 60 min, followed by immunoprecipitation with anti-FLAG antibodies and immunoblotting with anti-FLAG (left panel) or anti-HA (right panel) antibodies. Strains lacking either MAR1-FLAG or HAP2-HA were controls; input (represents ∼2.6% of the cell equivalents loaded in the eluate lane) for the *mar1;MAR1-FLAG;hap2;HAP2-HA(-)* sample is shown on the right panel.

### Biochemical and functional assays demonstrate an association between MAR1 and fusogen HAP2

To test for a biochemical interaction between these temporally co-functioning, co-localized proteins, we immunoprecipitated MAR1-FLAG from the lysates of activated *minus* gametes expressing both MAR1-FLAG and HAP2-HA and used immunoblotting to assess whether HAP2-HA was also present in the immunoprecipitates. As shown in Fig. 4E, HAP2-HA was indeed present in the anti-FLAG immunoprecipitates, but absent in control immunoprecipitates from activated *hap2::HAP2-HA(-)* gametes (lacking MAR1-FLAG), and activated *mar1; MAR1-FLAG(-)* gametes (lacking HAP2-HA). Notably, we consistently found that MAR1 preferentially associated with the upper, membrane-surface-expressed isoform of HAP2 (Fig. 4E). Despite their association, MAR1-FLAG expression (Fig. 5A) and localization (Fig. 5B) in *hap2(-)* gametes were indistinguishable from that of *minus* gametes expressing WT HAP2-HA. Furthermore, as reported here (Fig. 3D) and previously (Feng et al., 2018; Liu *et al.*, 2008), *hap2(-)* gametes were fully capable of undergoing mating structure adhesion with *plus* gametes, indicating that MAR1 was also functionally competent without HAP2.

**Figure 5.**
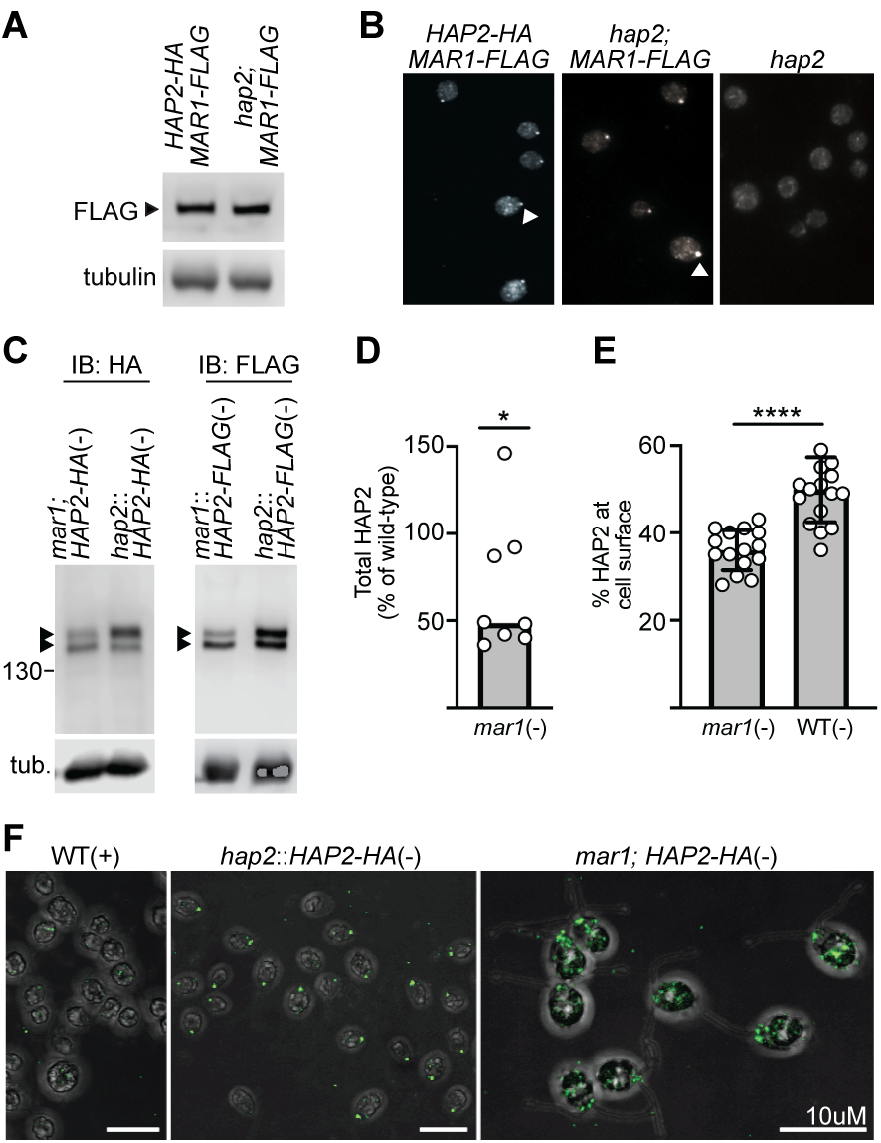
HAP2 expression and localization depend on MAR1, but MAR1 properties are independent of HAP2. (A-B) MAR1 expression and localization are independent of HAP2. **(A)** Immunoblots of MAR1-FLAG (upper panel) in lysates of *mar1;MAR1-FLAG;hap2;HAP2-HA(-)* gametes (left side) and *mar1;MAR1-FLAG;hap2(-)* gametes (right side). Tubulin is shown as a loading control (lower panel). **(B)** MAR1-FLAG immunolocalization in *hap2::HAP2-HA; MAR1-FLAG(-)* gametes (left), and *hap2::MAR1-FLAG(-)* gametes (middle) by fluorescence microscopy. Negative control (right), was *hap2(-)* gametes expressing neither HAP2-HA nor MAR1-FLAG. **(C-F)** HAP2 expression and mating structure localization are impaired in *minus* gametes lacking MAR1. **(C)** Representative immunoblots from lysates of HAP2-HA (left panel) or HAP2-FLAG (right panel) expressing *minus* gametes in the absence(left lanes) or presence (right lanes) of WT *MAR1*. Controls (right lanes) were *hap2::HAP2-HA(-)* and *hap2::HAP2-FLAG(-)* gametes which contained the endogenous WT *MAR1* gene; lower panel shows tubulin (tub) loading controls. **(D)** Quantification of total HAP2-HA and HAP2-FLAG in immunoblots that was normalized to tubulin for each sample. Median HAP2 expression (bar) in *mar1(-)* gametes was 49% of that in *minus* gametes expressing WT *MAR1*; Wilcoxon Signed Rank test, \**P*=0.04. Each open circle represents a biological replicate *mar1(-)* sample (*n*=8) and is expressed as a percentage of the median signal present in WT *MAR1(-)* samples (*n=*5). **(E)** Quantification of the percentage of total HAP2 present in the upper isoform of the HAP2 doublet that was determined from immunoblot signals of HAP2-HA or HAP2-FLAG expressed in *mar1(-)* or WT *MAR1(-)* gametes. The mean percent of cell surface HAP2 in *mar1(-)* gametes was 36% (*n*=15) compared to 50% in WT *MAR1(-)* gametes (*n*=17); two-tailed unpaired t test with Welch’s correction; \*\*\*\**P*<0.0001; bars are mean; error bars, SD. **(F)** Confocal z-stack composite images of anti-HA immunostained *mar1;HAP2-HA(-)* gametes (lacking *MAR1*, right), *hap2::HAP2-HA(-)* gametes (containing WT *MAR1*, middle), and control WT*(+)* gametes (left).

On the other hand, the expression and localization of HAP2 in *minus* gametes lacking MAR1 were substantially altered. Total HAP2 protein expression in *mar1;HAP2-HA(-)* and *mar1::HAP2-FLAG(-)* gametes was reduced 2-fold compared to that of strains expressing HAP2 in a WT *MAR1* background (representative blots, Fig. 5C and Supplemental Fig. S8; quantification of multiple independent experiments, Fig. 5D). Moreover, quantitative densitometry measurements of immunoblots across multiple biological replicates also showed that the mean percentage of HAP2 present in the upper, surface-expressed isoform was reduced from 50% in WT gametes to 36% in the *mar1(-)* gametes, representing a reduction of 28% (Fig. 5E; Supplemental Fig. S9).

Absence of MAR1 also substantially altered HAP2 localization. Immunofluorescent staining and confocal microscopy of HAP2-HA in *minus* gametes containing WT MAR1 showed that the predominant localization of HAP2-HA was, as shown above (Fig. 4C) and previously (Fedry *et al.*, 2017; Feng *et al.*, 2018; Liu *et al.*, 2010), as a single spot at the site of the *minus* mating structure (Fig. 5F, middle panel, Supplemental Fig. S9). Low intensity, non-specific anti-HA staining was observed in negative control WT*(+)* gamete samples (Fig. 5F, left panel), as well as in small amounts within cells and elsewhere on the slide in other samples (as shown above in Figs. 4C, 5B). On the other hand, HAP2-HA localization in *minus* gametes lacking MAR1 was strikingly different, and only rarely detected as a single apically-localized spot. Often, HAP2-HA in the *mar1(-)* gametes appeared as smaller puncta close to the site of the mating structure, but was predominately found throughout the cell body as multiple, discrete puncta of varying sizes (Fig. 5F, right panel, Supplemental Fig. S9). Thus, proper expression and localization of HAP2-HA at the *minus* mating structure depended upon MAR1.

## Discussion

We investigated the proteins and protein interactions that underlie membrane adhesion and bilayer merger during the gamete membrane fusion reaction in *Chlamydomonas*. We determined that the ectodomain of FUS1, the previously identified adhesion protein on the *plus* gamete mating structure, is predicted to be constituted entirely by a linear array of Ig-like domains, a domain architecture that BLAST searches and homology modeling show is similar to that of GEX2, a gamete adhesion protein present throughout land plants. We also identified a lineage-restricted, *minus* gamete-specific membrane protein, MAR1, that is localized at the surface of the *minus* mating structure and that binds directly to FUS1. Like FUS1, MAR1 is essential for mating structure adhesion and for gamete fusion, and these interacting proteins are now the second gamete membrane adhesion receptor pair demonstrated to be essential for fertilization (Bianchi *et al.*, 2014). In addition to interacting with FUS1, MAR1 is also associated with HAP2 on *minus* gametes and essential for proper HAP2 expression and localization during gametogenesis. Our results suggest that the membrane adhesion immediately preceding HAP2-dependent bilayer merger during fertilization in organisms across Viridiplantae is mediated by an interaction between a member of the conserved FUS1/GEX2 family of adhesion proteins and a lineage-restricted binding partner on the cognate gamete.

### FUS1/GEX2 joins HAP2 and KAR5/GEX1 to make a trio of protein families central to sexual reproduction in organisms from unicellular green algae to flowering plants

The Immunoglobulin Superfamily (IgSF) is large and diverse, and many of its members function in intercellular adhesion in prokaryotes and eukaryotes (Chatterjee et al., 2020; Honig and Shapiro, 2020). Several proteins important for sperm-egg interactions in metazoans possess Ig-like domains (Nishimura and L’Hernault, 2016), the best characterized being the sperm adhesion protein IZUMO1, whose single Ig-like domain forms a portion of the interface between IZUMO1 and its egg binding partner, JUNO (Aydin *et al.*, 2016; Ohto *et al.*, 2016). The previously reported presence of Ig-like domains in *Chlamydomonas* FUS1 (Misamore *et al.*, 2003) and plant GEX2 (Mori *et al.*, 2014) proteins called further attention to the widespread use these domains in sexual reproduction. Our findings here indicate, however, that the Ig-like domains of FUS1/GEX2 adhesion proteins constitute the entire functional portion of these proteins. The expansive binding repertoire of vertebrate antibody proteins is provided by structural variations in a subset of the immunoglobulin domains that entirely constitute them. Future structural studies should provide insights into whether similar variations in particular Ig-like domains of FUS1/GEX2 family members underlie their ability to interact with diverse gamete receptor proteins across taxa.

Remarkably, uncovering the breadth of conservation within this family of gamete adhesion proteins now indicates that at least three of the core functions unique to eukaryotic sexual reproduction are carried out by three protein families conserved across all plant lineages: (1) gamete membrane adhesion mediated by FUS1/GEX2 family members, (2) membrane merger mediated by HAP2 family members, and (3) pronuclear fusion mediated by the KAR5/GEX1 family of nuclear envelope proteins (Mori *et al.*, 2014; Ning *et al.*, 2013; Wong and Johnson, 2010). The HAP2 and KAR5/GEX1 families are not restricted to plant lineages and are essential for sexual reproduction in organisms from green algae and protists to multicellular plants and animals. The conservation of domain architectures within members of each of these three protein families suggests that while evolutionary pressures often lead to rapid changes in the primary amino acid sequence of reproductive proteins, this diversification might take place on a relatively unchanging structural backbone (Ferris et al., 1997; Mori *et al.*, 2014; Swanson and Vacquier, 2002).

### MAR1 is bifunctional and interacts with FUS1 on *plus* gametes and HAP2 on *minus* gametes during the membrane fusion reaction

Our biochemical results with endogenous proteins in gamete lysates and with bacterially expressed proteins indicated that interaction between FUS1 and MAR1 ectodomains was independent of *Chlamydomonas*-specific post-translational modifications and that it was direct. One potential binding site in the MAR1 ectodomain could be within its proline-rich region, which contains several PPSPX repeats akin to the poly-proline repeats of algal and higher plant hydroxyproline-rich glycoproteins (HPRGs, Ferris *et al.*, 2005; Ferris *et al.*, 2001) that support other types of protein-protein interactions. The MAR1 growth factor receptor domain, which is similar to growth factor motifs found in other gamete interaction proteins such as the mating type proteins in the ciliate, *T. thermophila* (Cervantes *et al.*, 2013); and Spe-9 in *C. elegans* (Singson *et al.*, 1998), could also be a potential protein interaction site.

Furthermore, results from several independent methods were all consistent with a biochemical and functional association between MAR1 and the class II fusogen HAP2 on the *minus* mating structure. Total HAP2 expression as well as the portion of HAP2 at the cell-surface were reduced in *minus* gametes lacking MAR1, HAP2 was mis-localized in gametes lacking MAR1, and MAR1 was biochemically associated with the cell-surface form of HAP2. One interpretation of these findings is that, like the partner protein interactors of viral class II fusion proteins, MAR1 participates in biosynthesis and trafficking of HAP2 to the *minus* mating structure. For example, in the absence of their partner E2 proteins, the E1 fusogens of Sindbis virus and Semliki Forest Virus are inefficiently transported to the plasma membranes of their host cells (Andersson et al., 1997; Carleton et al., 1997).

It will be important to determine whether the MAR1-HAP2 association is direct or indirect and to identify the regions of MAR1 and HAP2 that participate in this association. If the interaction is indirect, a handful of *Chlamydomonas* membrane proteins whose transcripts show *minus* gamete-specific expression similar to MAR1 (Ning *et al.*, 2013) would be good candidates for a linking protein; including Cre03.g175926, which has some sequence similarity to MAR1. If the interaction is direct, possible cytoplasmic domain interaction sites within HAP2 and MAR1 are a 237-residue segment of HAP2 essential for its mating structure localization (Liu et al., 2015) and a 400-residue segment within MAR1 with a predicted structural likeness to collagen (Supplemental Fig. S4).

Before a structure was available, Wong et al. (2010) demonstrated that the fusion capacity of *Arabidopsis* HAP2 was retained when its ectodomain was exchanged with the HAP2 ectodomain from a closely related species, but was not when the swap was with the ectodomain of a more distantly related species, indicating that the ectodomain of HAP2 was important in lineage-specific functions. Recent protein structures of the *Chlamydomonas* trimeric HAP2 ectodomain have brought to light regions within loops of domain I that may participate in such lineage-specific protein-protein interactions (Baquero et al., 2019; Feng *et al.*, 2018). Furthermore, comparisons between the HAP2 structures of *Chlamydomonas*, *Arabidopsis*, and *Trypanosoma cruzi* showed substantial, taxa-specific differences in their fusion loops, supporting a possible lineage-specific role for protein interactions of this region (Fedry et al., 2018).

Our discovery that lineage-specific MAR1 functions at the nexus of two broadly conserved, fusion-essential protein families leads to the speculation that the cognate gamete adhesion receptor pairs in other organisms might also be composed of one conserved protein and another that is lineage specific. The IZUMO1-JUNO receptor pair would support such a model, as JUNO is specific to mammals, but IZUMO1 is present throughout vertebrates (Bianchi *et al.*, 2014; Grayson, 2015). Even this model is overly simplistic, however, since the FUS1/GEX2 family member in *Chlamydomonas* is present in the gamete lacking HAP2, whereas the FUS1/GEX2 family members in land plants are present in the gamete expressing HAP2.

Nevertheless, our findings raise the exciting possibility that MAR1 binding to *Chlamydomonas* FUS1 triggers conformational changes within HAP2 required for its fusogenic function. In class I fusogen-dependent Paramyxoviruses, the interaction of a fusogen-associated adhesion protein with its membrane receptor, activates the fusion protein (Navaratnarajah et al., 2020). Future studies on the FUS1-MAR1-HAP2 axis have the potential to illuminate new, conserved regions in HAP2 that act before its function in bilayer merger in organisms across plant taxa as well as in the many unicellular organisms and metazoans that also depend on HAP2 for gamete fusion.

## Acknowledgments

We thank UMD colleagues Drs. Jun Zhang, Mayanka Awasthi, and Peeyush Ranjan; UT Southwestern colleagues Drs. Muqing Cao, Wenhao Li, Jue Ning, and Saikat Mukhopadhyay; Institut Pasteur colleagues Drs. Felix Rey, Eduard Salazar, and Ignacio Fernandez; and colleague, Ursula Goodenough, Washington University, St. Louis, for helpful discussions; Dr. Tim Maugel of the UMD Laboratory for Biological Ultrastructure for assistance with SEM; and Drs. Kate Luby-Phelps and Abhijit Bugde (UT Southwestern Medical Center, Live Cell Imaging Core) and Amy Beaven (UMD, Imaging Core) for guidance with light microscopy. Funding was provided by NIH GM56778 and GM122565 to W.J.S. and F32-GM126735 to J.F.P.

## Author Contributions

J.F.P., Y.L., and W.J.S. designed the experiments. J.F.P. and Y. L. performed the experiments. J.F.P., Y.L., and W.J.S. analyzed the results and prepared the manuscript.

## Declaration of Interests

The authors declare no competing interests.

## Materials and Methods

### Cells and cell culture

*Chlamydomonas* strains *21gr(+)*, *fus1(+), fus1::FUS1-HA(+), hap2(-)*, *hap2::HAP2-HA(-)*, *CC-5313(+)*, *mar1(-)* also called *LMJ.RY0402.185187*, and *CC-5325(-)* were used for experiments. New strains used in this study were made by stable transformation and crosses of the above strains and are fully described in Supplemental Fig. S7 and S8, along with the *Chlamydomonas* Resource Center codes for previously-generated strains. Cells were grown vegetatively in liquid TAP medium under a 13:11 hr light-dark cycle at 22°C. Gametogenesis was induced by transferring vegetative cells into N-free medium followed by overnight agitation on a shaker in continuous light.

### *Chlamydomonas* transformation and crosses

*Chlamydomonas* cells were transformed by electroporation of plasmid DNA encoding the tagged HAP2 or MAR1 transgenes. Supplemental Figs. S6, S8, Table S1, and Supplemental Methods provide details for how plasmid construction, cell transformation, and identification of transformants based on growth on selective media, PCR screening, and expression of tagged proteins were performed. For crosses (Supplemental Fig. S7), equal numbers of the indicated *plus* and *minus* gametes were mixed together and plated onto paromomycin (15 μg/mL, Sigma) TAP-agar plates at 0.5, 1, 1.5, and 2 hours after mixing. Supplemental Methods describes the techniques used for zygote maturation, induction of meiosis, and selection of colonies.

### Immunoprecipitations, protease treatments, and recombinant protein production

Activated *hap2(-)* gametes surface biotinylated with Sulfo-NHS-LC-Biotin (Pierce) were mixed with an equal number of *FUS1-HA(+)* gametes or, as a control, with *fus1(+)* gametes. The samples were lysed, FUS1-HA partners were affinity purified using anti-HA beads, bound proteins were eluted with HA peptide, and biotinylated proteins were detected on immunoblots using streptavidin-conjugated HRP. The 250 kDa regions of similarly prepared samples were analyzed by mass spectrometry (Supplemental Fig. S1 and Supplemental methods). Protease treatments were performed as described previously (Liu *et al.*, 2010) with minor modifications (Supplemental Methods).

For protein association assays, lysates were prepared from the indicated strains after gamete activation by mixing gametes for 30-60 min with either fusion-defective *fus1(+)* or *hap2(-)* gametes. The mixed gametes were disrupted by lysis in detergent buffers, centrifuged, and the supernatants used for immunoprecipitation with anti-FLAG or anti-HA antibodies (Supplemental Methods). The immunoprecipitates on protein A agarose or sepharose were either boiled directly in sample buffer or eluted with FLAG peptide (Sigma, 200 μg/mL) or HA peptide (Roche, 1 mg/mL) for detection by immunoblotting. The presence of tagged forms of HAP2 and MAR1 proteins in immunoprecipitation eluates and gamete lysates was assessed by SDS-PAGE followed by immunoblotting as described in the Supplemental Methods. Details of recombinant protein production in *E. coli* and pull-down experiments are also described in the Supplemental Methods. Immunoblots were developed with SuperSignal West Femto (Thermo Scientific) and exposed to film or scanned with a C-digit blot scanner (LI-COR). Images were processed using Image Studio Digits software (LI-COR) and Adobe Photoshop.

### Microscopy

Ciliary adhesion between *minus* and *plus* gametes was assessed by differential interference contrast (DIC) microscopy and scanning electron microscopy (SEM, Supplemental Methods). Immunofluorescent staining of gametes was performed as described previously (Belzile *et al.*, 2013; Feng *et al.*, 2018) with some modifications (Supplemental Methods). Images were processed with Adobe photoshop software. Images from the Leica SP5 X confocal are z-stack composites made with Leica LASAF software.

### Bioassays

Assays used to quantify gamete activation, mating structure adhesion and cell-cell fusion were performed as previously described (Feng *et al.*, 2018) with minor modification (Supplemental Methods). For assessment of gamete activation, at least 3 independent cell wall loss experiments were performed for each cross tested. For adhesion and fusion experiments, at least 200 cells (as either singles or pairs) were counted in each of at least 5 biological replicates per cross.

### MAR1 polyclonal antibody generation

Antibodies against MAR1 peptides (1 = TQPPRPPWPPRPPPAPPPS, residues 164-182; and 2 = QIPQAPRWPYPQLPSWPPAS, residues 214-233) were generated in rabbits by YenZym Antibodies (Brisbane, CA). Only polyclonal antibodies generated against peptide 1 were used. Antibodies were purified on peptide-conjugated affinity columns and specificity verified by immunoblotting with recombinant protein samples with and without MAR1 (Supplemental Fig. S6).

### Quantification and Statistical Analyses

Data are presented as median or mean ± SD and analyzed using Prism 9.0 software (GraphPad). The statistical tests used to analyze data from each experiment are indicated in the figure legends. Surface and total tagged HAP2 protein signals on digital images of immunoblots were quantified using Image Studio Digits software (LI-COR) and tubulin was used for normalization as described in the Supplemental Methods.

## *Chlamydomonas* transformation

*Chlamydomonas* cells were transformed by electroporation of plasmid DNA (Shimogawara et al., 1998) using a BioRad GenePulser Xcell. *Spe*I-linearized *MAR1-FLAG* or *Sbf*I-linearized *HAP2-FLAG* plasmid DNA were transformed into *hap2(-)* and *mar1(-)* cells. The sequences of the plasmid DNAs used for transformation and of PCR products across the insertion site of the *mar1(-)* CLiP mutant, *LMJ.RY0402.185187,* were confirmed by analysis at the MGH CCIB DNA core and Eurofins, respectively. Positive transformants were selected for further analysis based on their growth on zeocin (10 μg/mL, Invitrogen) TAP-agar plates, presence of the *ble* gene product assessed by PCR of Chelex-100-extracted template DNA (Nouemssi et al., 2020) using primers Zeo_F1 and Zeo_R1, and immunoblotting of gamete lysates for expression of the FLAG-tagged proteins.

## *Chlamydomonas* zygote maturation

Agar plates from crosses were exposed to continuous light for 16-24 hrs, then placed in the dark to allow zygote maturation. After at least 7 days in the dark, any remaining unfused cells were scraped towards the edges of the plate with a sterile razor blade and their further growth prevented by exposure of the plate to chloroform vapors for 30-60 sec. The thick cell walls of the zygospores remained attached to the agar and were resistant to the cytolytic effects of the chloroform vapors. After incubation under a 13:11 light-dark cycle at 21°C for 2 weeks to allow germination and visible colony growth, 20 – 30 colonies (each representing the vegetative haploid descendants of 4 initial meiotic progeny) were picked and resuspended in N-free medium with shaking for 1 hr to obtain a single-cell suspension, then re-plated onto TAP-agar plates to allow for growth of single colonies composed of individual meiotic progeny. Resulting colonies were screened for mating type, genotype and phenotype by PCR, western blotting, and behavior in mixing assays (Supplemental Fig. S7 and S8) to isolate the desired genotypes.

## Recombinant protein production and interaction assays

For recombinant protein production, the expression vectors GST-rFUS1 and HIS-rMAR1 plasmids were transformed into BL21(DE3) (New England Biolabs) competent *E. coli* cells. Bacterial cells expressing the recombinant proteins were grown with shaking overnight at 37°C in liquid LB media containing antibiotics and then 0.5 mL was inoculated into 50 mL LB cultures. The cultures were shaken for 1 hour at 37°C and induced with IPTG (1 mM) for 3 hours at 30°C. Bacteria were lysed in 0.9 mL lysis buffer (20 mM Tris pH 8.0, 150 mM NaCI, 5 mM DTT, 1 mg/mL lysozyme, 5 μg/mL DNase I, 10 μg/mL RNase A, and protease inhibitor cocktail) on ice for 30 min. Triton X-100 was added to 1% (final concentration), then samples were sonicated on ice for 1 min, cleared by centrifugation at 4°C 15,000 x g for 30 min and used in GST pull-down assays. *BL21/pGST-FUS1* supernatants were incubated with glutathione beads for 1 hour and washed 3 times with washing buffer (20 mM Tris, 150 mM NaCI, 0.3% Triton X-100, protease inhibitor cocktail). The glutathione beads with bound GST-FUS1 were incubated with *BL21/pHis-MAR1* lysate overnight at 4°C, washed 3 times with washing buffer, and incubated with elution buffer (50 mM Tris pH 8.0, 20 mM reduced glutathione, 0.3% Triton X-100, protease inhibitor cocktail). Recombinant GST protein was used as a control for the recombinant protein interaction assays.

## Surface Biotinylation

To prepare *hap2(-)* gametes for surface biotinylation, they were activated in db-cAMP buffer for 30 min (Sigma; Misamore et al., 2003) before a 30 min incubation in 1 mg/mL Sulfo-NHS-LC-Biotin (Pierce). Biotinylation was stopped by addition of 20 mM Tris (pH 7.3) followed by 3 washes in N-free medium. Biotinylated *hap2(-)* gametes were mixed with an equal number of *FUS1-HA(+)* gametes for 30 min followed by lysis in 4 mL RIPA buffer (20 mM Tris, 150 mM NaCI, 1% NP-40, 0.5% DOC, 0.1% SDS and protease inhibitor cocktail, Wilson et al., 1999) and sonication 3 times (10 sec each) on ice. The lysate was cleared by centrifugation for 30 min at 15,000 x g and incubated for 4 hrs with anti-HA monoclonal antibody (Santa Cruz) and protein A agarose (Sigma). The protein A agarose was washed 4 times with 1/3 RIPA buffer (20 mM Tris, 150 mM NaCI, 0.33% NP-40, 0.17% DOC, 0.03% SDS with protease inhibitor cocktail) and eluted with 1 mg/mL HA peptide as described previously (Ning et al., 2013). Biotinylated proteins in the eluate were detected on immunoblots using streptavidin-HRP (Jackson ImmunoResearch).

## Protease Treatments, Immunoprecipitations and Immunoblotting

To assess surface localization of HAP2-HA and HAP2-FLAG, live *minus* gametes were treated with 0.05% trypsin for 20 min followed by 3 washes in N-free media with 0.1 mg/mL Trypsin inhibitor from chicken egg white (Sigma) before lysis in SDS-PAGE sample buffer. Surface localization of MAR1-FLAG, HAP2-HA and HAP2-FLAG was also assessed by treating live gametes with 1% pronase (1.0 mg/mL, Sigma) for 20 min followed by 3 washes in N-free media with 1 mM PMSF before lysis in SDS-PAGE sample buffer.

The presence of tagged forms of HAP2 and MAR1 proteins in *minus* gamete lysates and immunoprecipitation eluates was detected by immunoblotting. For immunoprecipitations, the mixed gametes were disrupted by lysis in RIPA buffer or 1% Triton X-100 (20 mM Tris, 150 mM NaCI, 1% Triton X-100, protease inhibitor cocktail (Sigma or Roche) as indicated, followed by centrifugation. Anti-FLAG (Sigma, mouse M2 monoclonal or rabbit polyclonal) or anti-HA (Santa Cruz, mouse F-7 monoclonal) antibodies at a dilution of ∼1/100 in the lysates were used for immunoprecipitations.

For gamete lysates, 2.5 × 10^7^ naive gametes were suspended in 225 μL N-free medium, and lysed by adding 75 μL of 4x SDS-PAGE sample buffer (1× concentration in sample = 40 mM Tris, pH 6.8, 1% SDS, 5% glycerol, 0.0003% Bromophenol blue, 1 mM EDTA and either 100 mM DTT or 5 mM TCEP) at 95°C for 5-10 min. 2.5 × 10^6^ cell equivalents were subjected to electrophoresis on SDS polyacrylamide tris-glycine gels (7 or 8%), transferred to PVDF membranes, blocked in 5% milk + TBSt buffer, and probed with either rat anti-HA (1/1000; Sigma, 3F10) or mouse anti-FLAG (1/5000, Sigma, M2) primary antibodies, followed by goat-anti-rat HRP or goat-anti-mouse HRP (1/5000; Millipore) secondary antibodies diluted in 0.5% milk + TBSt buffer. Tubulin was used as a loading control and was detected by probing with mouse anti-*α*-tubulin primary antibodies (1/5000; Sigma, B512).

## Quantification of relative protein abundance

Median signal intensities were determined within rectangles drawn around the upper band of HAP2, the lower band of HAP2, and *α*-tubulin on digital images of immunoblots. Background correction was performed by ImageStudioDigits software (LI-COR) using a right/left border setting of 2 on each rectangle. Total HAP2 signal was calculated as the sum of the signals of the upper and lower HAP2 bands. To compare total HAP2 proportions respective to tubulin loading controls across multiple blots, lane normalization factors were calculated for both total HAP2 and *α*-tubulin as described by the vendor (LI-COR). The percentage of surface-expressed HAP2 for each sample was calculated as [(signal of HAP2 upper band)/(Total HAP2 signal)] ×100.

## Sequence homology detection and protein structure modeling

The similarity of the ectodomain sequences of *Chlamydomonas reinhardtii* FUS1 (GenBank#:AAC49416) to plant GEX2 species was detected by DELTA-BLAST (Boratyn et al., 2012) and PSI-BLAST (Altschul et al., 1997; Aravind and Koonin, 1999), with significant hits to GEX2 family members detected upon the 3^rd^ iterative search. The online tool PROMALS3D (Pei and Grishin, 2007) was used to construct a pairwise alignment between *reinhardtii* FUS1 and *Arabidopsis thaliana* GEX2 (DDBJ#:AB743888) ectodomain sequences (Supplemental Fig. S4). The RaptorX web server (Xu, 2019; Xu et al., 2020; Zeng et al., 2018) was used to construct structural models of the *C. reinhardtii* FUS1 and *A. thaliana* GEX2 ectodomains and to identify previously undetected Ig-like domains. Images of the FUS1 and GEX2 structural models (Fig. 1B) were generated using PyMol software (Schrodinger, LLC). Structural homologies of GEX2, FUS1 and MAR1 (GenBank: KT288268) with existing structures in the Protein Data Bank were detected by use of RaptorX (Källberg et al., 2012), PHYRE2 (Kelley et al., 2015), SWISS-MODEL (Waterhouse et al., 2018) and HHPRED (Zimmermann et al., 2018).

## Differential interference contrast and scanning electron microscopy

Ciliary adhesion between *minus* and *plus* gametes was assessed by differential interference contrast (DIC) microscopy using an inverted Axiovert 135 TV microscope fitted with an ORCA-ER (Hamamatsu) digital camera. For examination of ciliary adhesion by scanning electron microscopy, WT*(+)* gametes were mixed with *mar1;hap2(-)* gametes for 30 min, washed 10x in sterile-filtered N-free media, fixed for 20 min in Parducz fixative (1 part saturated HgCl_2_, 6 parts 2%OsO_4_)(Parducz, 1967), then aliquots of the cells suspended in fixative were transferred onto 0.1% poly-L-lysine coated coverslips for an additional 45 min. Coverslips were washed 10x in ddH2O, then dehydrated by sequential washes (3x each, 2 min/wash) in 70%, 95%, and 100% ethanol solutions. Coverslips were critical point dried from CO_2_ and mounted on stubs over double-sided tape before being sputter-coated with gold–palladium (60%:40%) (Balzers Med 010) and observed using a scanning electron microscope (Hitachi S-4700 FESEM) at an accelerating voltage of 5 kV in high vacuum.

## Assessing gamete activation, mating structure adhesion, and gamete fusion

Gamete activation as measured by cell wall loss was quantified by the amount of OD435-detectable chlorophyll released into the supernatant upon treatment with a buffer containing 0.075% Triton X-100 as described in (Snell, 1982; Wang et al., 2006). Mating structure adhesion was assayed as indicated previously (Feng et al., 2018) in gametes mixed at a concentration of 5.0×10^7^/mL, followed by fixation at 10 min after mixing with an equal volume of 5% glutaraldehyde. Ciliary adhesions were disrupted by pipetting 20x with a 200 μL pipette tip. Glutaraldehyde fixation coupled with the light agitation from pipetting disrupts flagellar adhesions leaving pairs of *plus* and *minus* gametes attached only by their mating structures (Feng *et al.*, 2018; Forest, 1983), The percent of cells attached by their mating structures was calculated as: [(2 × number of pairs) / (2 × number of pairs + number of single gametes)] × 100. Determination of gamete fusion was assessed by the microscopic enumeration of quadri-ciliated cells as previously described (Feng *et al.*, 2018). Gamete strains used for quantification of gamete activation, mating structure adhesion, and gamete fusion were the F_1_ and F_2_ progeny strains (Supplemental Fig. S8) derived from crosses (Supplemental Fig. S7B) of the original *mar1(-)* CLiP mutant strain *LMJ.RY0402.185187,* which had been transformed with the MAR1-FLAG encoding transgene (Supplemental Fig. S6). Preliminary results gathered using the *LMJ.RY0402.185187(-)* gametes agreed with the phenotypes observed in the *mar1(-)* progeny strains.

## Indirect immunofluorescence

Immunofluorescent staining of gametes was performed on samples of approximately 1 × 10^7^ live gametes in N-free medium that were allowed to settle and adhere for 30 min on 0.1% poly-L-lysine coated slides. Excess media and non-adhered cells were gently removed by pipetting before slides were plunged into ice-cold methanol for 20 min. For single staining with anti-HA antibodies (1/100, Sigma, 3F10), gametes were blocked for 30 min in diluted goat serum (5% goat serum, 0.1% BSA, 1% cold-water fish gelatin in PBS). For double staining with anti-HA (1/100, Sigma, 3F10) and anti-FLAG antibodies (1/100, Sigma, M2), gametes were blocked in diluted donkey serum (5% donkey serum, 0.1% BSA, 1% cold-water fish gelatin in PBS). Primary antibodies were diluted in blocking sera and incubated with samples for 2 hrs at room temperature, washed 3 times with PBS, and incubated with secondary antibodies for 2 hrs at room temperature. For single staining with anti-HA, the secondary antibody used was Alexa Fluor 488-conjugated goat anti-rat IgG (1/400, Invitrogen); for double staining, the secondary antibodies were Alexa Fluor 488 Donkey Anti-Rat and Alexa Fluor 594 Donkey Anti-Mouse (Invitrogen). Samples were then washed 3x with PBS and coverslips were mounted using Fluoromount-G (Southern Biotech). Immunofluorescence images of gametes were captured on either an Axioplan2 or III RS fluorescent microscope in FITC or Rhodamine channels as described previously (Misamore *et al.*, 2003) or a Leica SP5 X Laser Scanning confocal microscope with a Leica 63x/1.4 NA oil objective lens as indicated.

---

**Supplemental Figure Legend S1.**
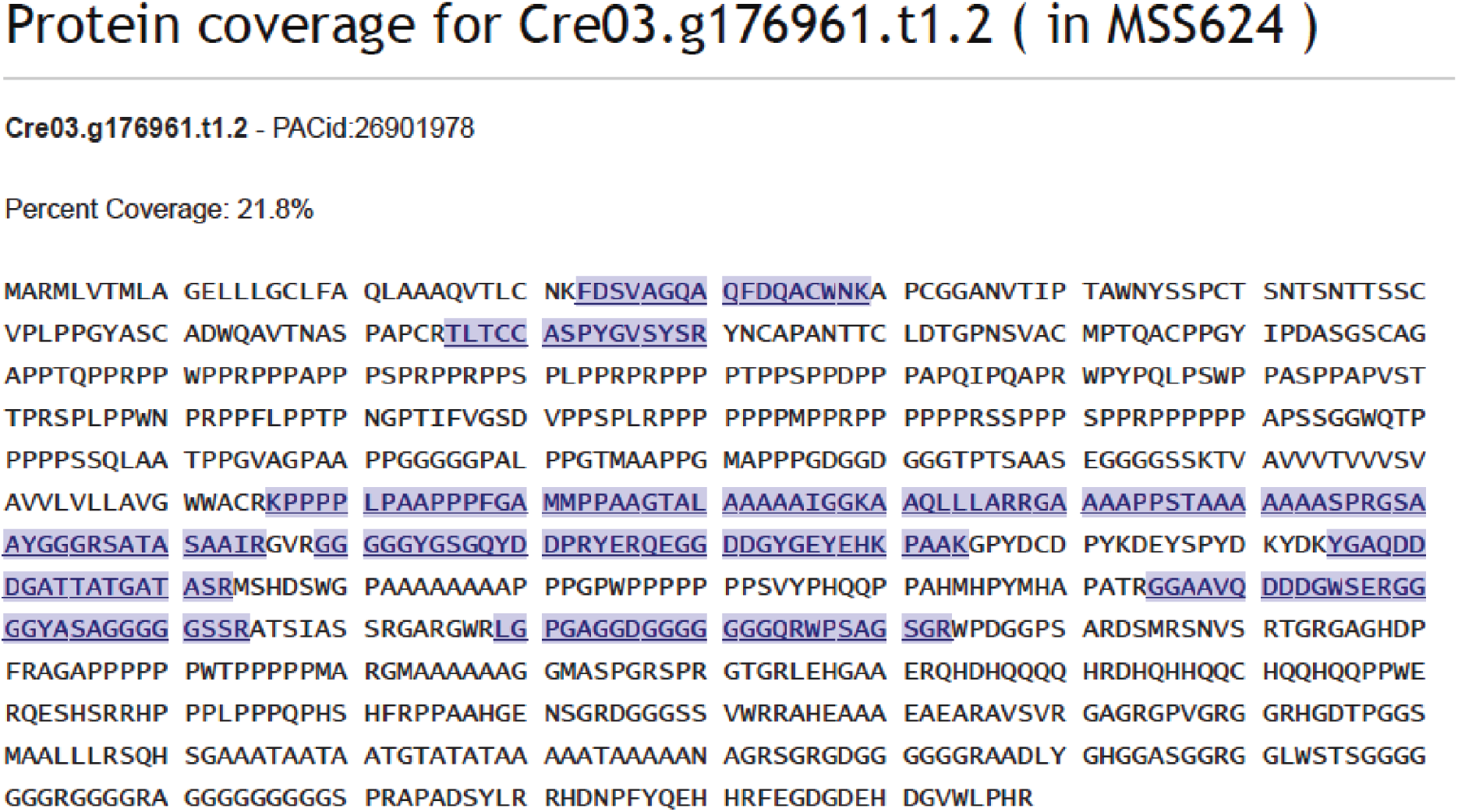
The streptavidin-staining protein of ∼250 kDa indicated by the arrow in Fig. 1C is Cre03.g176961. The region corresponding to the streptavidin-stained band on the immunoblot shown in Fig. 1C was excised from an SDS-PAGE gel identical to the one used for the immunoblot (except that elution was done by boiling instead of by HA-peptide) and sent for mass spectrometry analysis by the Proteomics Core at UT Southwestern Medical Center (Dallas, TX). The Cre03.g176961 peptides identified by mass spectrometry in the sample are underlined and highlighted blue.

**Supplemental Figure Legend S2.**
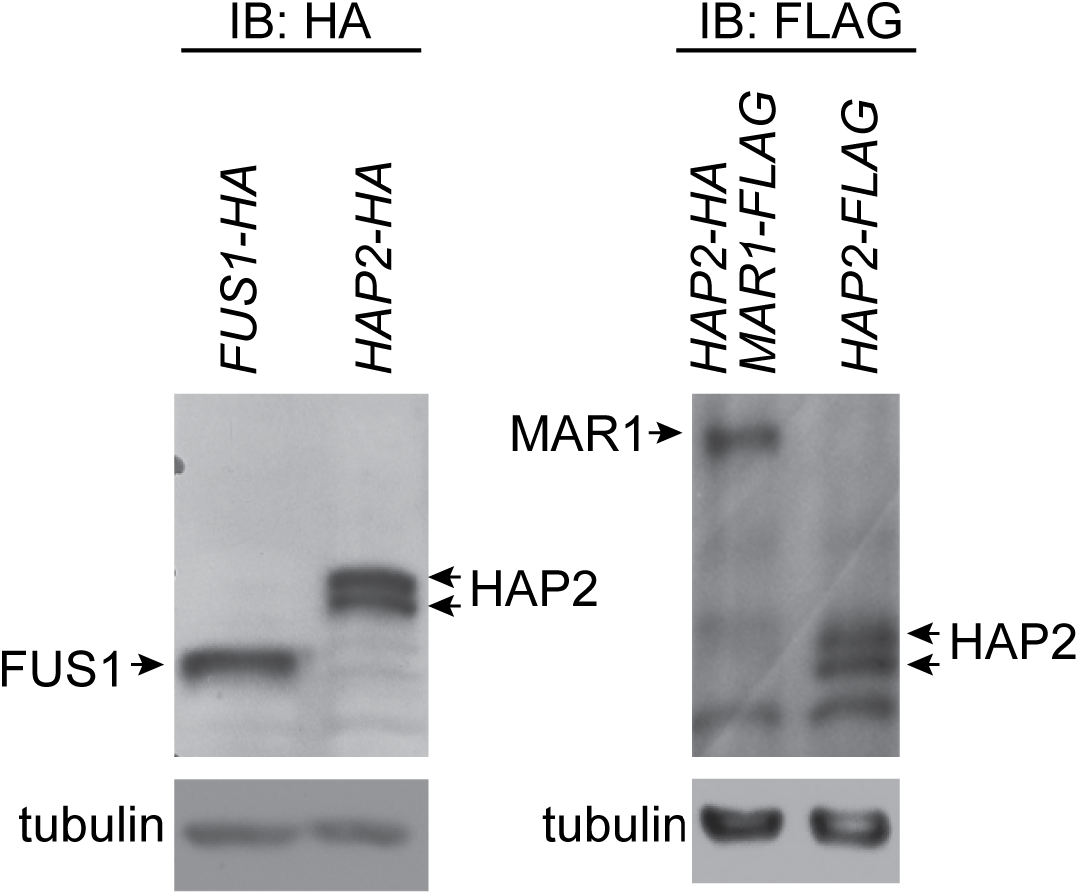
In spite of differences in transcript levels (Fig. 1D), immunoblotting of *plus* and *minus* gametes indicate that the protein levels of HAP2, MAR1 and FUS1 are similar. Lysates from equal numbers of naive *fus1*::*FUS1-HA(+)*, *hap2::HAP2-HA(-)*, *hap2::MAR1-FLAG;HAP2-HA(-),* and *hap2*::*HAP2-FLAG(-)* gametes were analyzed by immunoblotting with anti-HA or anti-FLAG antibodies. The tubulin blots document equivalent loading. Proteins of interest are labeled.

**Supplemental Figure Legend S3.**
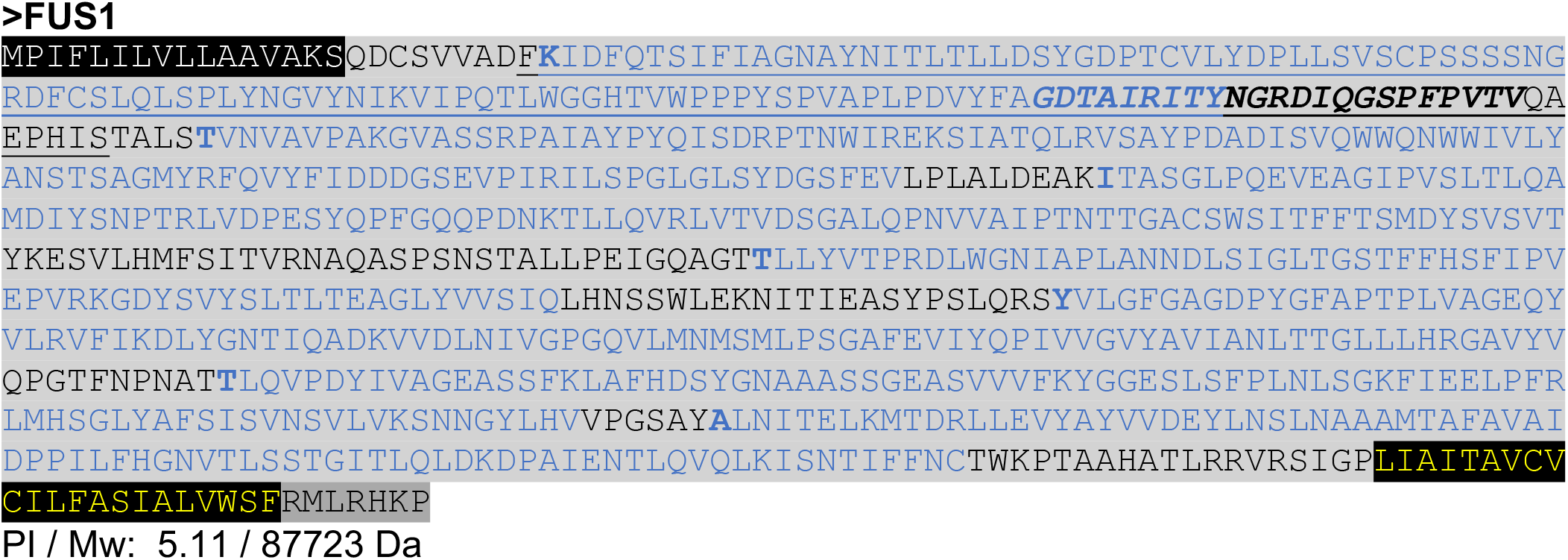
Annotated amino acid sequence of FUS1. GenBank: AAC49416. Signal peptide called by SignalP (Nielsen, 2017). Extracellular region: <u>Immunoglobulin-like fold domain</u> (IPR013783, CATH 2.60.40.10) called by InterProScan (Mitchell et al., 2019), and seven Ig-like domains predicted by structural homology models. Only the predicted beta sheet portion of each non-canonical Ig-like domain is in blue text with the first amino acid of each bolded. ***Filamin/ABP280 repeat-like domain*** (IPR017868) called by InterProScan. Transmembrane domain called by Phobius (Käll et al., 2007). Intracellular region. FUS1 has no predicted intrinsically disordered regions (Necci et al., 2017) and no regions of amino acid enrichment (MotifScan). The theoretical isoelectric point (pI) and molecular weight (Mw) were computed without signal peptides using the Expasy tool Compute pI/Mw. Amino acid sequence similarities between FUS1 and bacterial invasin proteins were previously reported by Misamore *et al.* (2003). Partial structural homology of the *Chlamydomonas* FUS1 ectodomain with bacterial proteins was found with HHPRED (S-layer protein; PDB: 5FTX, prob: 99.83%, E-value 4e-15, score: 169.78, ident: 14%) and with an invasin-like protein structure; FdeC, from *E. coli* (PDB: 4E9L, prob: 99.7%, E-value: 2.2e-13, score: 147.25, ident: 17%); PHYRE2 ranked the gelation factor rod domain from *Dictyostelium* as the top hit (PDB 1WLH, confidence: 98.7%, ident: 17%) closely followed by the invasin-like protein FdeC (PDB 4E9L, residues 381-620, confidence: 97.3, ident: 16%); and Swiss-Model also found bacterial templates from invasin-like and S-layer protein structures as top hits (PDB: 4E9L, coverage: 45%, ident: 19.4%; PDB: 5FTX, coverage: 49%, ident: 12.4%). Sequence-based searches using BLAST for FUS1-related proteins found the following algal orthologs: BBC28482.1 in *Eudorina sp*.; BAU61607.1 in *Gonium pectoral*e; BBC28430.1 in *Yamagishiella unicocca*; and KAF5841765.1 in *Dunaliella salina*. The *Arabidopsis* GEX2 ectodomain was also found to have partial structural homology with the same bacterial proteins and *Dictyostelium* gelation factor rod domain that showed structural similarity to FUS1. HHPRED showed GEX2 homology with the *Dictyostelium* gelation factor (PDB 1WLH, prob: 99.7%, E-value 9.5e-14, score: 149.33, ident: 14%) the invasin-like protein structure; FdeC, from *E. coli* (PDB: 4E9L, prob: 99.6%, E-value: 4.6e-11, score: 134.9, ident: 13%) and the S-Layer protein (PDB: 5FTX, prob: 99.3%, E-value: 9.3e-8, score: 114.9, ident: 11%). PHYRE2 ranked the *Dictyostelium* gelation factor rod domain as the top hit for GEX2 (PDB 1WLH, confidence: 99.8%, ident: 21%), followed by hits to filamin and cadherin-1 structures before the invasin-like protein FdeC (PDB 4E9L, confidence: 99.2, ident: 14%). Swiss-Model also found the *Dictyostelium* gelation factor as the top hit for GEX2 (PDB: 1WLH, coverage: 33%, ident: 17.3%) followed hits to filamin structures before the invasin-like and S-Layer protein structures (PDB: 3TKV -superseded by 4E9L, coverage: 44%, ident: 16.5%; PDB: 5FTX, coverage: 24%, ident: 15.5%).

**Supplemental Figure Legend S4.**
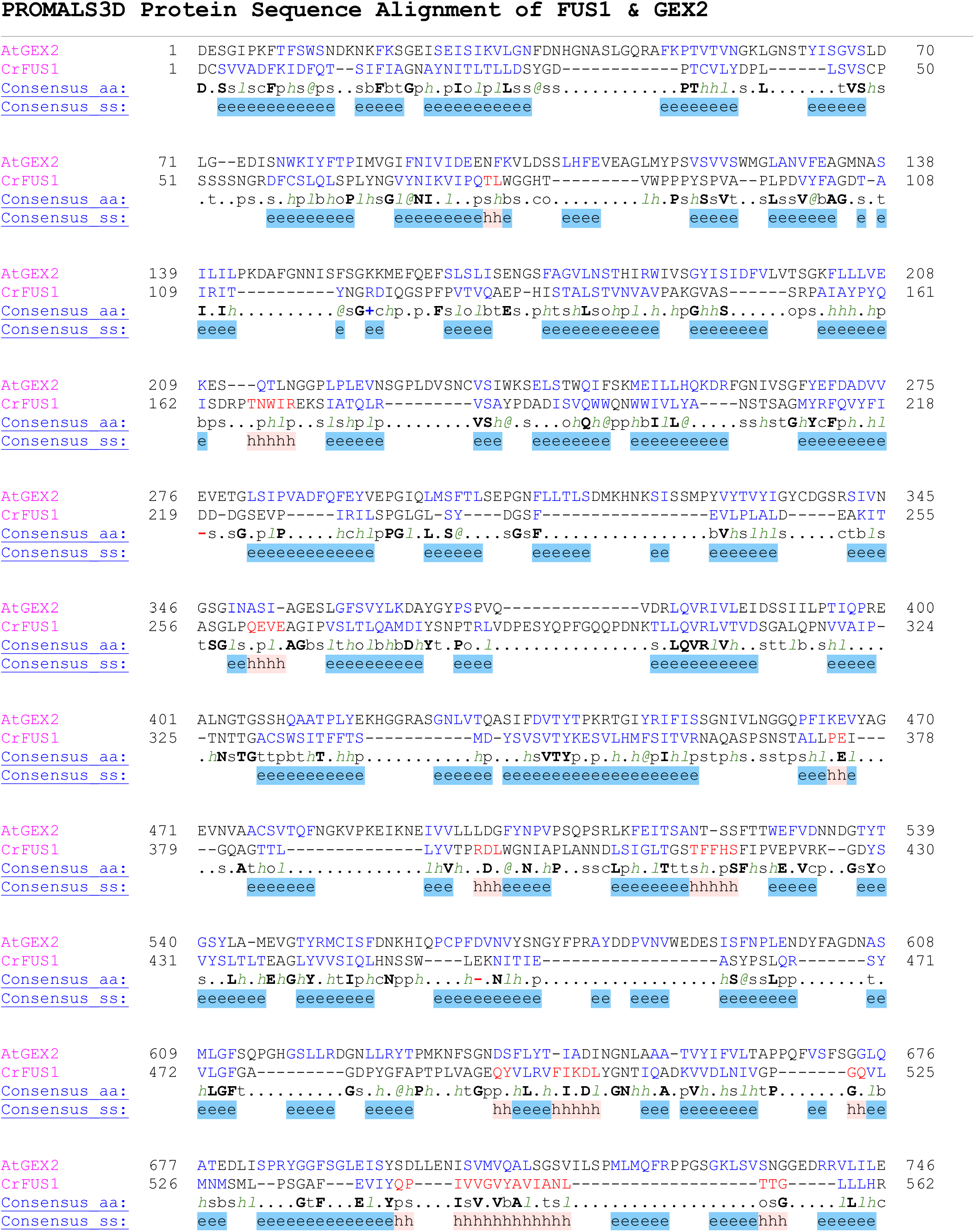
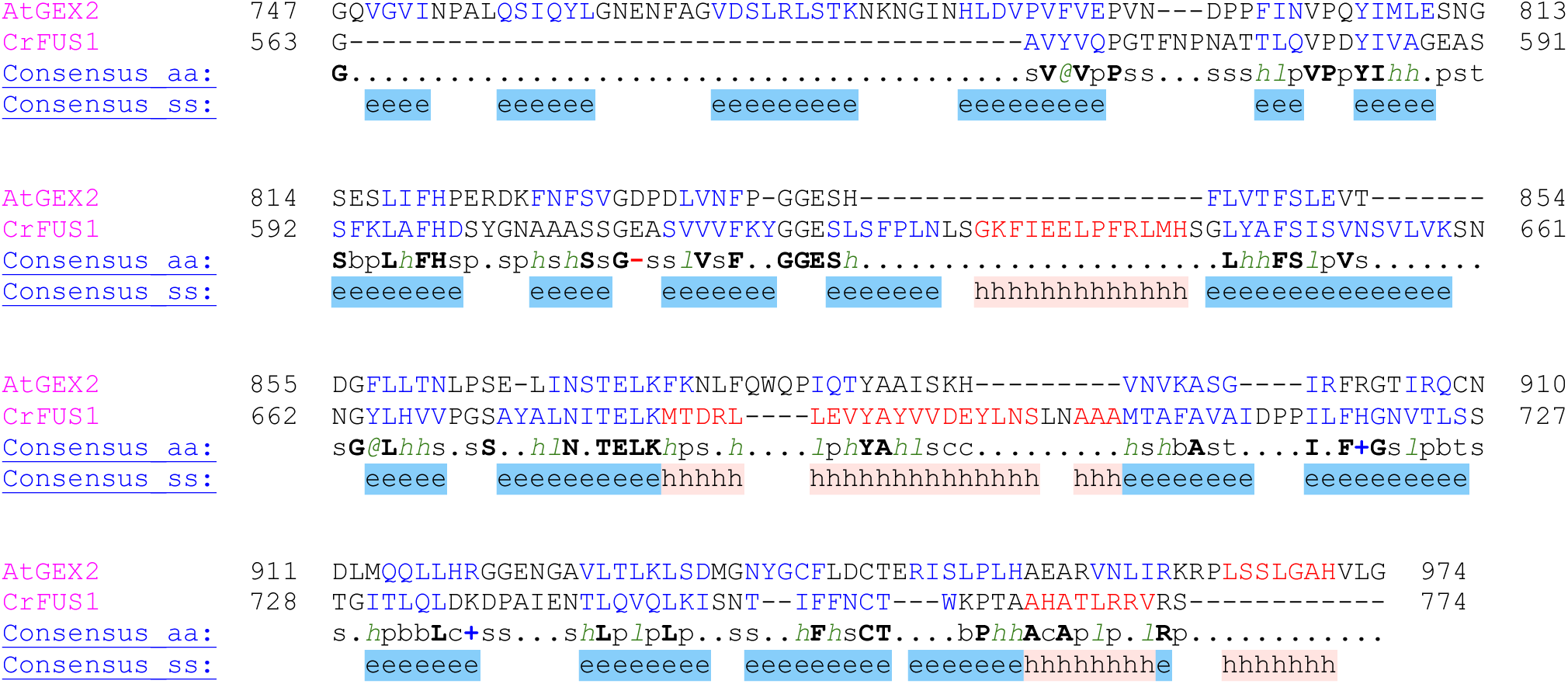
FUS1-GEX2 alignment. A PROMALS3D (Pei and Grishin, 2007) amino acid alignment of *Chlamydomonas reinhardtii* FUS1 (CrFUS1) (GenBank#:AAC49416, residues 18-791) and *Arabidopsis thaliana* GEX2 (AtGEX2) (DDBJ#:AB743888, residues 28-1001) ectodomain sequences with consensus predicted secondary structure below (consensus_ss, PSI-PRED, Jones, 1999). The CrFUS1 ectodomain sequence was 15.9% identical to the AtGEX2 sequence, and about ¾ as long (72.3% coverage). Predicted secondary structure is color coded onto the residues of each sequence and shown as a consensus below the alignment as: beta strand (`e`) and alpha helix (`h`). For consensus_aa residue code: conserved (bold & uppercase); aliphatic (*l*); aromatic (*@*); hydrophobic (*h*); alcohol (o); polar (p); tiny (t); small (s); bulky (b); positively charged (+); negatively charged (-); charged (c).

**Supplemental Figure Legend S5.**
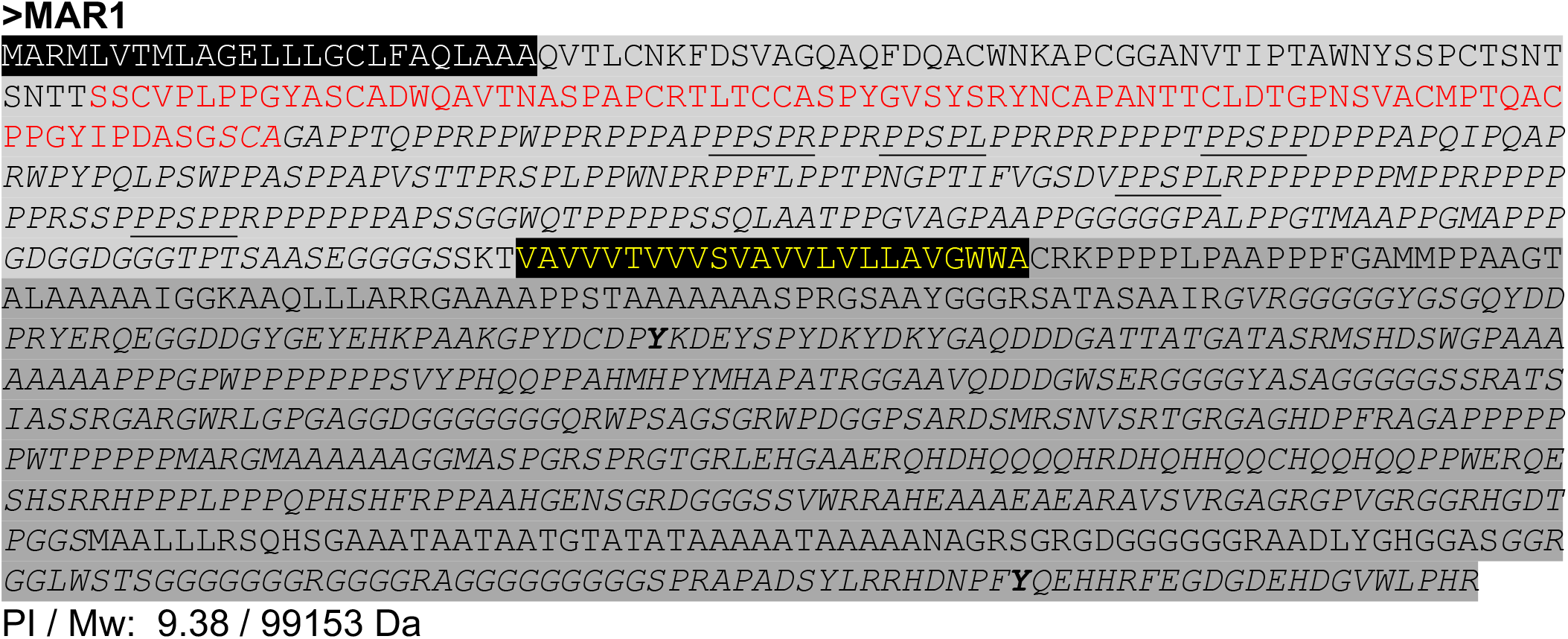
Annotated amino acid sequence of MAR1. Signal peptide called by SignalP (Nielsen, 2017). Extracellular region. Growth factor receptor cysteine-rich domain superfamily (IPR009030) called by InterProScan. (**Y**) Putative tyrosine kinase phosphorylation sites called by both NetPhos3.1 (Blom et al., 2004) and MotifScan (Pagni et al., 2007). *Disordered regions* called by MobiDB-lite (Necci *et al.*, 2017). The following amino acids are enriched in the indicated sequence spans (MotifScan): Ala (A) 423-493; 845-921(15.1%) / Gln (Q) 773-802(3.6%) / Gly (G) 334-385; 625-979(15.6%) / His (H) 767-828(3.0%) **/** Pro (P) 142-365; 590-621; 726-738(19.1%). In the proline-rich (33% prolines) ectodomain, there are 5 repeats of a <u>PPSPX</u> motif found in other *Chlamydomonas* Hydroxy-Proline Rich Glycoproteins (HRGP). Transmembrane domain called by Phobius (Käll *et al.*, 2007). Theoretical isoelectric point (pI) and molecular weight (Mw) were computed without signal peptide using Expasy tool Compute pI/Mw. Structural homology modeling of MAR1 offered insights into the intracellular region. RaptorX template-based models of residues 512-620 were based on an ABA receptor kinase protein structure from *Arabidopsis* (PDB: 5XD6, score:78, p-value: 5.77e-07) and included prediction of a staurosporine binding site (pocket multiplicity of 52). PHYRE2 models of residues 619-1018 were based on a collagen template structure (PDB: 1YGV, confidence: 99.7%, identity: 15%). Sequence-based searches using BLAST for MAR1-related proteins found the following putative homologs: *Vocar.0015s0371* and *Vocar.0015s0370*, in *Volvox carteri; Cre03.g175926* in *Chlamydomonas*, and Genbank: KXZ56873.1 in *Gonium pectorale*.

**Supplemental Figure Legend S6.**
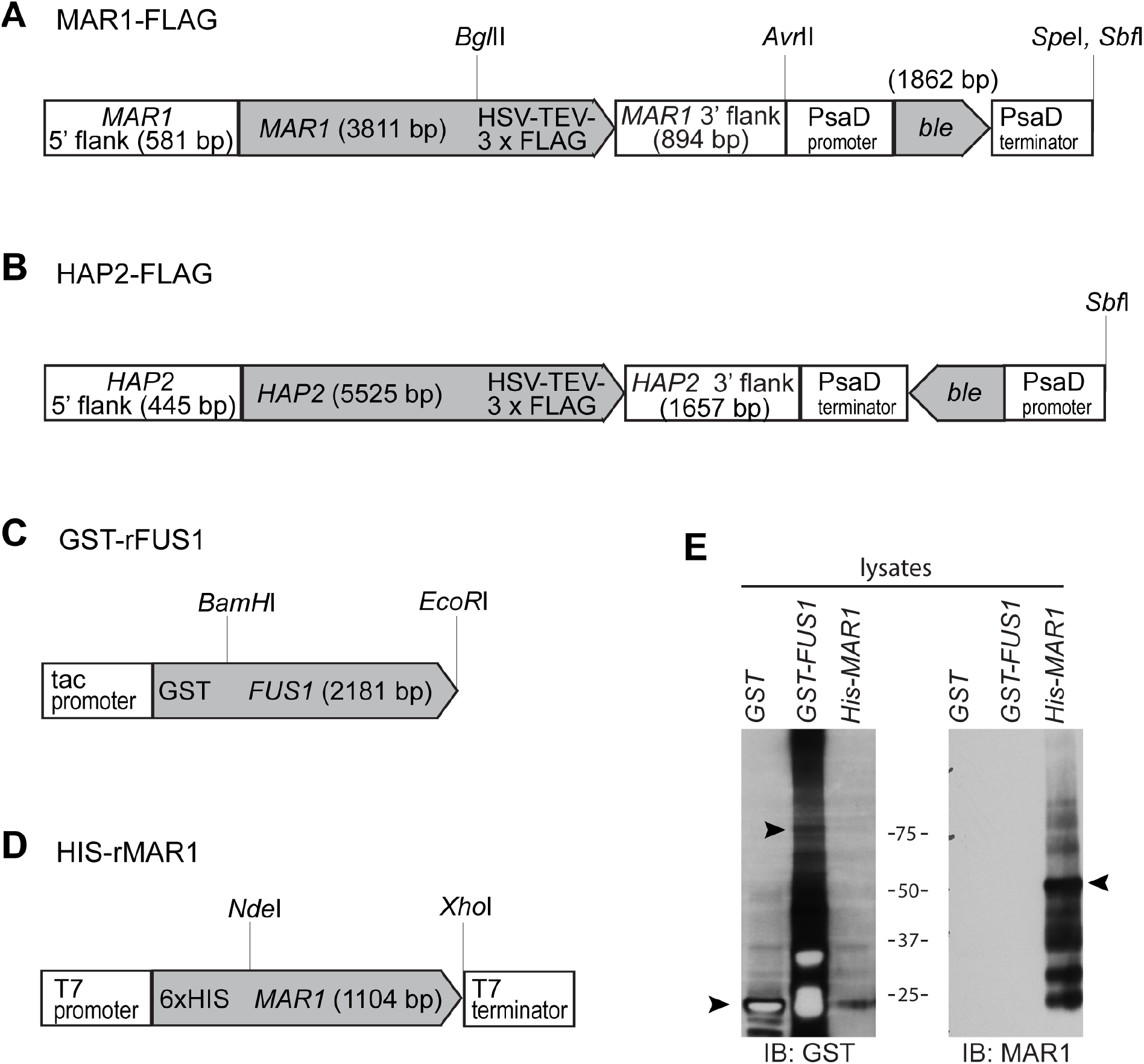
Construction of plasmids and specificity tests of the anti-MAR1 polyclonal antibody on recombinant proteins (Yenzyme). **(A)** The *MAR1-FLAG* plasmid was made by amplifying the *MAR1* gene (Cre03.g176961, 5981 bp) including its 581 bp 5’ flanking promoter region and 1589 bp 3’ flanking region using primers L8 and R5B, genomic DNA from *minus* strain *B215* as template, and the GC-rich PCR system (Roche). The PCR product was restricted at an *Avr*II site to truncate the 3’ flanking termination region to 894 bp, and cloned into a *pGEM-T-easy* vector (Promega). An oligonucleotide encoding an HSV-TEV-3 x FLAG epitope tag peptide (IDT) was added to the C-terminus of the *MAR1* gene using standard PCR weaving methods. The zeocin-resistance-encoding *ble* gene was inserted into the *MAR1-FLAG* plasmid at *Avr*II/*Spe*I sites using an In-Fusion Dry-Down PCR Cloning Kit (Takara). Primers L20 and G176961.R20 were used for amplification of the *ble* CDS and *PsaD* promoter and termination regions (1862 bp) using *pGenD-Ble* plasmid DNA as template; *Chlamydomonas* Resource Center (Fischer and Rochaix, 2001). **(B)** The *HAP2-FLAG* plasmid was made by inserting an oligonucleotide encoding tandemly arranged HSV-TEV-3 x FLAG epitope tag into the C-terminus of the existing *HAP2* plasmid pYJ36 (Liu et al., 2008) using standard PCR weaving methods. The *ble* gene was inserted into the *HAP2-FLAG* plasmid at a *Spe*I & *Xba*I site using an Infusion Dry-down PCR Cloning Kit (Clontech). Primers PGenD.s and PGenD.as were used for amplification of the *ble* CDS and *PsaD* promoter and termination regions using *pGenD-Ble* plasmid DNA as template. **(C)** The *GST-rFUS1* plasmid used to express the recombinant GST-tagged FUS1 ectodomain in B21 bacteria (Qiagen) was a pre-existing expression vector (Misamore *et al.*, 2003) containing an oligonucleotide encoding residues 17 – 747 of the FUS1 ectodomain in a *pGEX-2T* vector (Amp^+^) backbone. **(D)** The *HIS-rMAR1* plasmid used to express the recombinant His-tagged MAR1 ectodomain protein in BL21 bacteria was made by first amplifying an oligonucleotide encoding MAR1 residues 1 - 417 from *MAR1* cDNA using primers g176961-L3 and g176961-R10. This PCR product was cloned into a T-easy vector (Promega), and then a segment encoding residues 26 to 389 of the MAR1 ectodomain was further cloned by GenScript into the PET28a(+) vector (Kan^+^) using *Nde*I & *Xho*I sites. **(E)** MAR1 antibody (Yenzyme) specificity test in lysates of induced *BL21/pGST, BL21/pHis-rMAR1* and *BL21/pGST-rFUS1* bacterial cells were blotted with MAR1 and GST antibody. Arrowheads show the recombinant MAR1, FUS1 and GST proteins.

**Supplemental Table S1.**
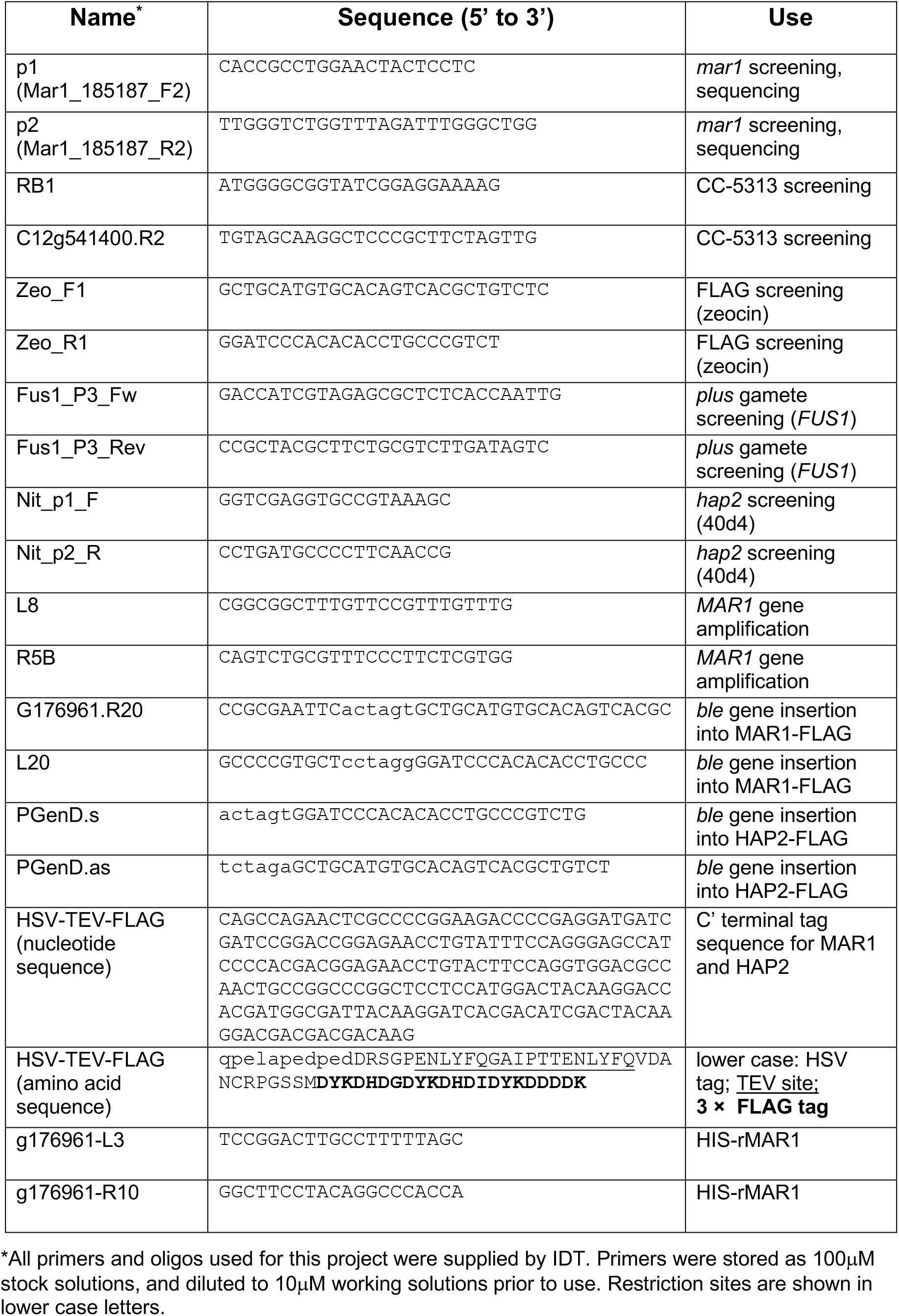

**Supplemental Figure Legend S7.**
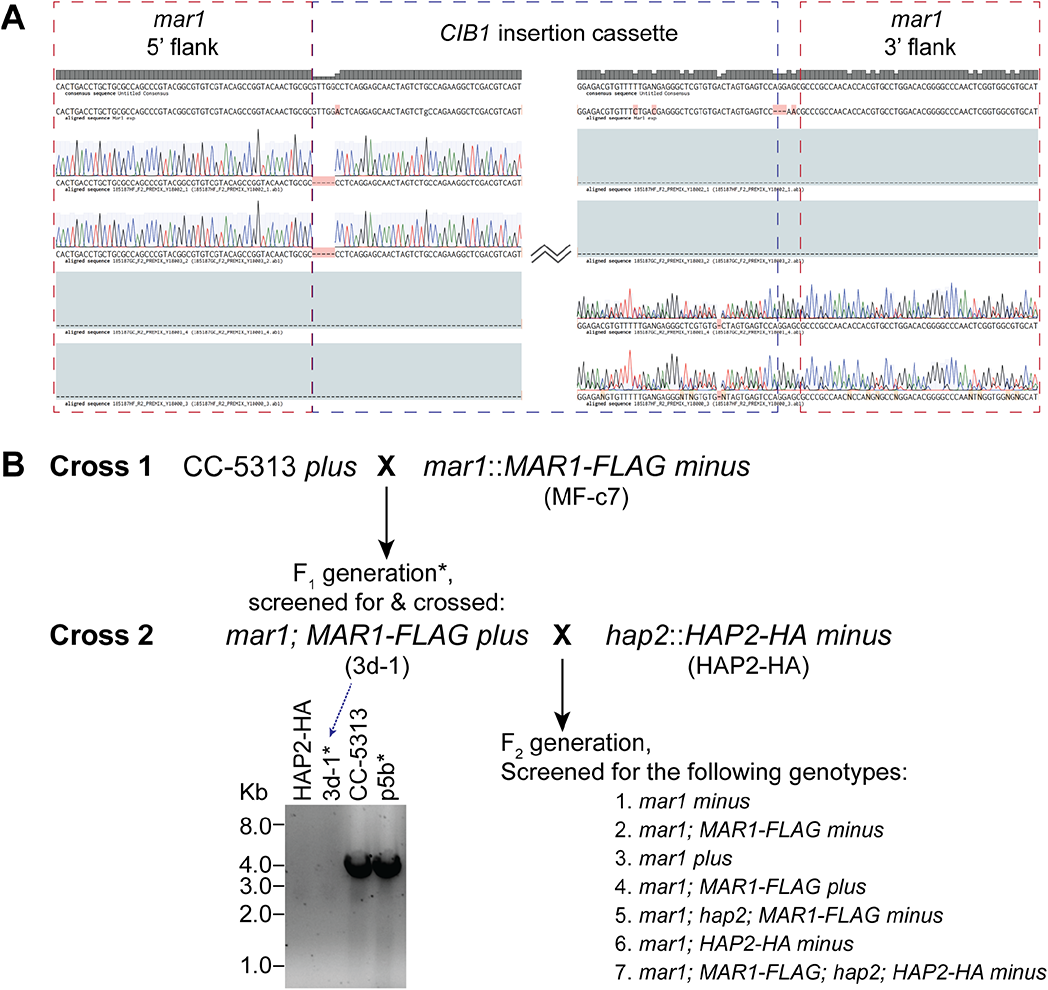
*mar1* insertion site and genetic crosses**. (A)** Aligned sequence traces from across the *CIB1* insertion site of the *mar1(-)* deletion strain (CLiP mutant *LMJ.RY0402.185187*). PCR products amplified using *mar1(-)* genomic DNA as template were purified and single tube sequenced by Eurofins with the same primers (p1 and p2). A Benchling (San Francisco, CA) alignment of sequencing results shows the *mar1(-)* sequence directly flanking the CIB1 insertion site. The *CIB1* cassette inserted in the antisense direction at the location expected by its CLiP library annotation (Li et al., 2019). At the site of insertion, the junction of the *CIB1* cassette and *mar1* 5’ flank lacked an expected “*GTTGGA*”; and the junction of the *CIB1* cassette with the *mar1* 3’ flank contained a *“GG”* insertion. **(B)** Schematic depicting the *Chlamydomonas* genetic crosses performed. Resulting progeny were analyzed 1) for co-segregation of the *mar1* mutant genotype with the fusion-defective phenotype in *minus* gametes, and 2) to obtain specific genotypes in the F_2_ generation (bottom right). The *mar1(-)* strain used in Cross 1 was transformed with *MAR1-FLAG* in order to obtain F_1_ progeny. The *CC-5313(+)* gamete strain used in Cross 1 had a lesion in the gene Cre12g541400 that was intentionally not selected for (to avoid possible confounding effects) in the *3d-1(+)* gamete F_1_ strain used for Cross 2. Screening for the transmission of the *CC-5313* lesion by DNA gel electrophoresis of PCR products using primers RB1 and C12g541400.R2 is shown on the bottom left, with the 4013bp product indicating the presence of the insertional lesion. In the F_2_ generation, single progeny colonies were picked and screened for the presence or absence of genetic and phenotypic markers (Supplemental Fig. S8) to isolate the indicated genotypes (bottom right).

**Supplemental Figure Legend S8.**
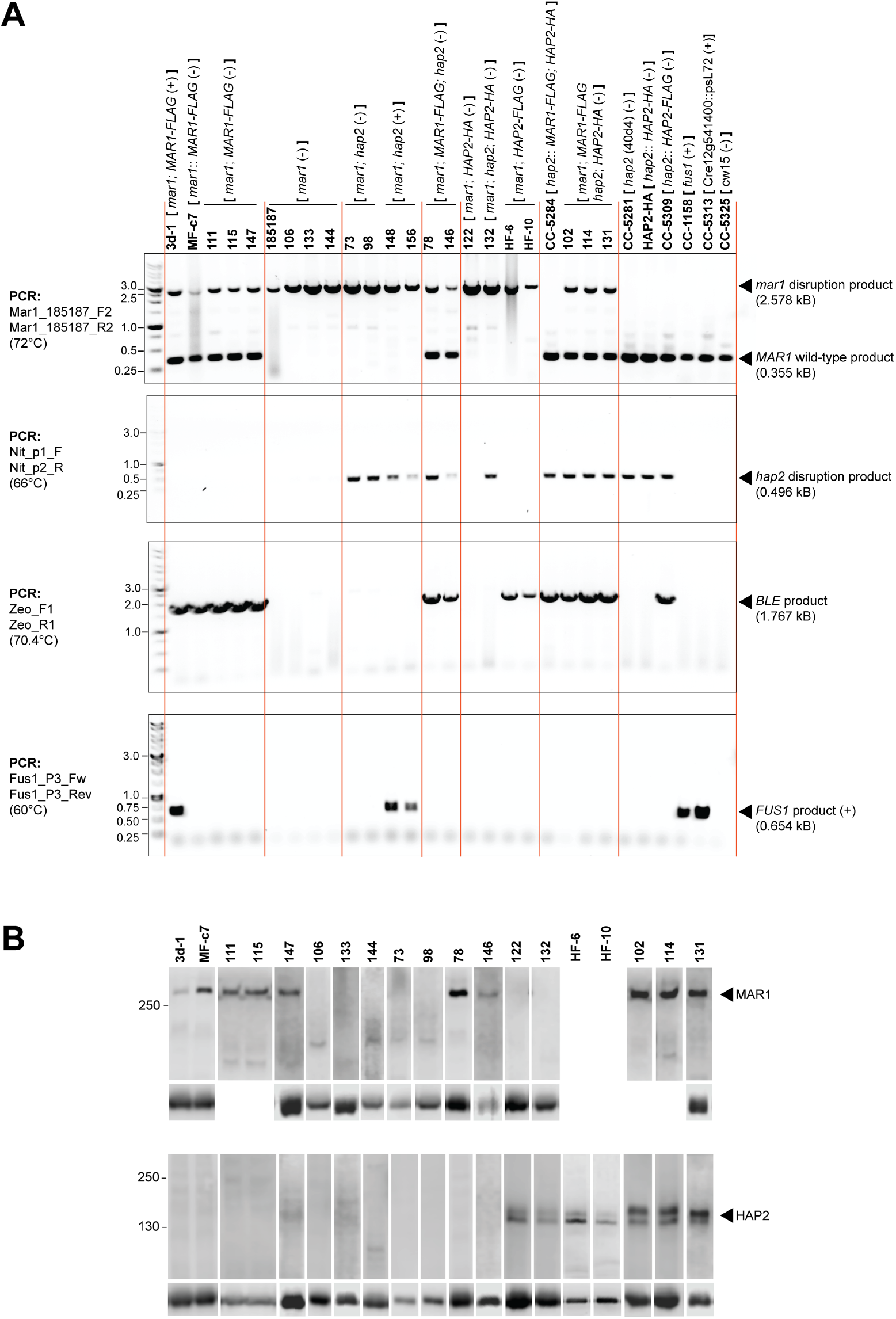
Genotypic and phenotypic screening results of strains used in this study. **(A)** Images of DNA gel electrophoresis results from PCR screening of selected *Chlamydomonas* progeny obtained in crosses (Supplemental Fig. S7). Genotypes are shown in the brackets above each strain number (top). Strains previously deposited in the *Chlamydomonas* Resource Center are identified by their CC numbers. Ladder (kB), primers used in PCR, and annealing temperatures are shown on the left. Expected PCR product sizes (kB) and their identities are shown on the right. **(B)** Immunoblots showing the presence or absence of MAR1-FLAG, HAP2-HA and HAP2-FLAG proteins in naive gamete lysates from the strain number indicated (top) and tubulin loading control (below). For HAP2 blots, HAP2-FLAG was detected with anti-FLAG antibodies for samples from strains HF-6 and HF-10; HAP2-HA was detected with anti-HA antibodies for all other sample blots. For MAR1 blots, MAR1-FLAG was detected with anti-FLAG antibodies. Blots for 111, 115, 102, and 114 were stripped and re-probed and the corresponding tubulin signal is shown below the HAP2 blot. Samples from *mar1*::*HAP2-FLAG(-)* strains were not screened for MAR1-FLAG expression. *Chlamydomonas* Resource Center codes and citations for other strains used in this study that have been previously described are: • *21gr(+);* CC-1690, used as a wild-type (+) strain (Proschold et al., 2005) • *hap2(-);* CC-5281, the 40D4 mutant (Liu et al., 2015) • *hap2::HAP2-HA(-);* CC-5295 (Feng *et al.*, 2018; Liu *et al.*, 2015) • *hap2::HAP2-FLAG(-);* CC-5309, in the 40D4 mutant background (Liu et al., 2010) • *fus1(+);* CC-1158 (Ferris et al., 1996) • *fus1*::*FUS1-HA*(+); referred to in text and shown in Fig. 2B and C simply as “*FUS1-HA*” (Liu *et al.*, 2010). • *CC-5313(+)*; used in genetic crosses • *CC-5325(-)*; the parental strain to the *mar1(-)* mutant (CLiP: LMJ.RY0402.185187, Li *et al.*, 2019)

**Supplemental Figure Legend S9.**
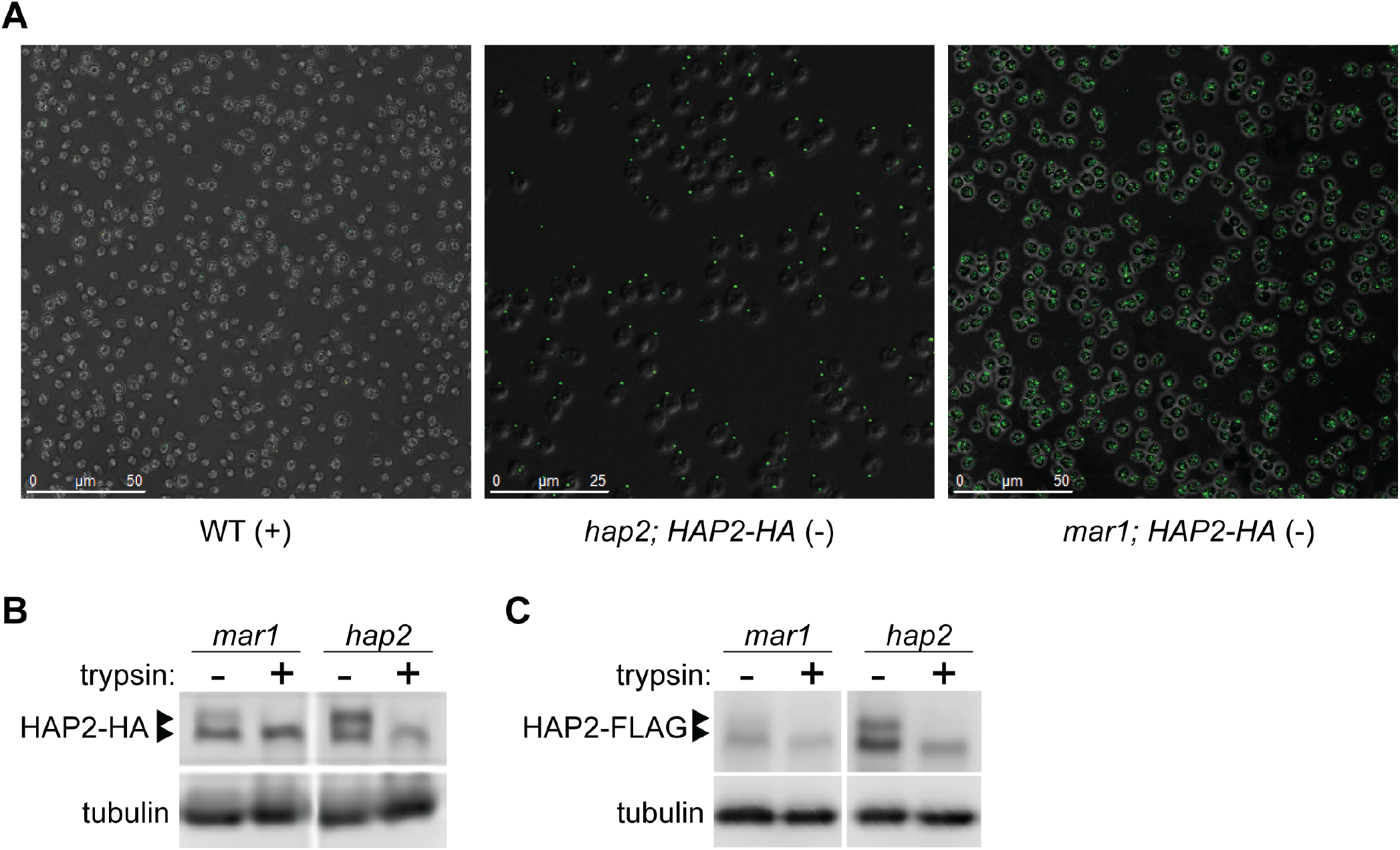
Expression of HAP2-HA in *mar1(-)* gametes. **(A)** Confirming results in Fig. 5F, wide field views of many gametes in merged bright field immunofluorescent confocal z-stack images show the broad mis-localization of HAP2-HA in *mar1(-)* gametes compared to the mating structure localization of HAP2-HA in positive control *hap2::HAP2-HA(-)* gametes. Negative control WT*(+)* gametes (*21gr*) are shown on the left. **(B)** Trypsin treatment of live *mar1;HAP2-HA(-)* gametes and **(C)** of live *mar1::HAP2-FLAG(-)* gametes confirmed that the minimal amounts of the surface-expressed HAP2 isoform (upper band) in gametes lacking *MAR1* was present at the cell surface. Immunoblots of trypsin-treated control samples of *hap2::HAP2-HA(-)* and *hap2::HAP2-FLAG(-)* gametes are shown in the right panels of (B) and (C).

## Notes

### Competing Interest Statement

The authors have declared no competing interest.

